# The Deceptively Simple N170 Reflects Network Information Processing Mechanisms Involving Visual Feature Coding and Transfer Across Hemispheres

**DOI:** 10.1101/044065

**Authors:** Robin A. A. Ince, Kasia Jaworska, Joachim Gross, Stefano Panzeri, Nicola J. van Rijsbergen, Guillaume A. Rousselet, Philippe G. Schyns

## Abstract

A key to understanding visual cognition is to determine *where*, *when* and *how* brain responses reflect the processing of the specific visual features that modulate categorization behavior—the *what.* The N170 is the earliest Event-Related Potential (ERP) that preferentially responds to faces. Here, we demonstrate that a paradigmatic shift is necessary to interpret the N170 as the product of an information processing network that dynamically codes and transfers face features across hemispheres, rather than as a local stimulus-driven event. Reverse-correlation methods coupled with information-theoretic analyses revealed that visibility of the eyes influences face detection behavior. The N170 initially reflects coding of the behaviorally relevant eye contralateral to the sensor, followed by a causal communication of the other eye from the other hemisphere. These findings demonstrate that the deceptively simple N170 ERP hides a complex network information processing mechanism involving initial coding and subsequent cross-hemispheric transfer of visual features.

## Introduction

The ultimate goal of cognitive neuroscience is to understand the brain as an organ of information processing. We subscribe to the assumption that the information processing systems of the brain, like all information processing systems, can be fruitfully described at different levels of abstraction, with specific contributions from different levels of granularity of brain signals (Marr, 1982; Tanenbaum and Austin, 2012). However, analysis at any level of abstraction will remain difficult unless we understand more directly *what* information the brain processes when it categorizes the external world. For example, our brain can quickly detect the presence of a face, implying that brain networks can extract and process the specific visual information required for face detection. As experimenters, we typically do not have a detailed description of such task-specific information and so we cannot explicitly test hypotheses about its algorithmic processing in brain signals.

Here, we address this issue by first isolating *what* specific information modulates face detection behavior. Then we examine, *where, when* and *how* this face information modulates dynamic signals of integrated brain activity on the left and right hemispheres. Since neural activity produces these integrated signals, from them we can derive the timing and approximate regions where neural populations are processing the specific face information underlying behavioral responses.

In humans, the N170 is the first integrated measure of cortical activity that preferentially responds to faces, with larger amplitudes to entire faces than to stimuli from other categories (Bentin et al., 1996; Rossion and Jacques, 2008). We developed this account, demonstrating that the N170 waveform reflects a feature coding mechanism (Schyns et al., 2003, 2007; Smith et al., 2004; van Rijsbergen and Schyns, 2009; Rousselet et al., 2014a). With face stimuli, coding starts with the eye contra-lateral to the recording sensor (e.g. the left eye on the right sensor, see Figure 1), on the downward slope of the N170 (~140 ms post-stimulus), followed in some face categorizations by the coding of task-relevant features—e.g. coding of the diagnostic contra-lateral wrinkled nose corner in ‘disgust,’ or the diagnostic corner of the wide-opened mouth in ‘happy’. By coding, we refer to the N170 time windows when the single-trial visibility of face features—randomly sampled with the Bubbles procedure, which uses a number of randomly positioned Gaussian apertures on individual trials to sample contiguous pixels from the face stimuli—covaries with the corresponding single-trial EEG responses.

**Figure 1.**
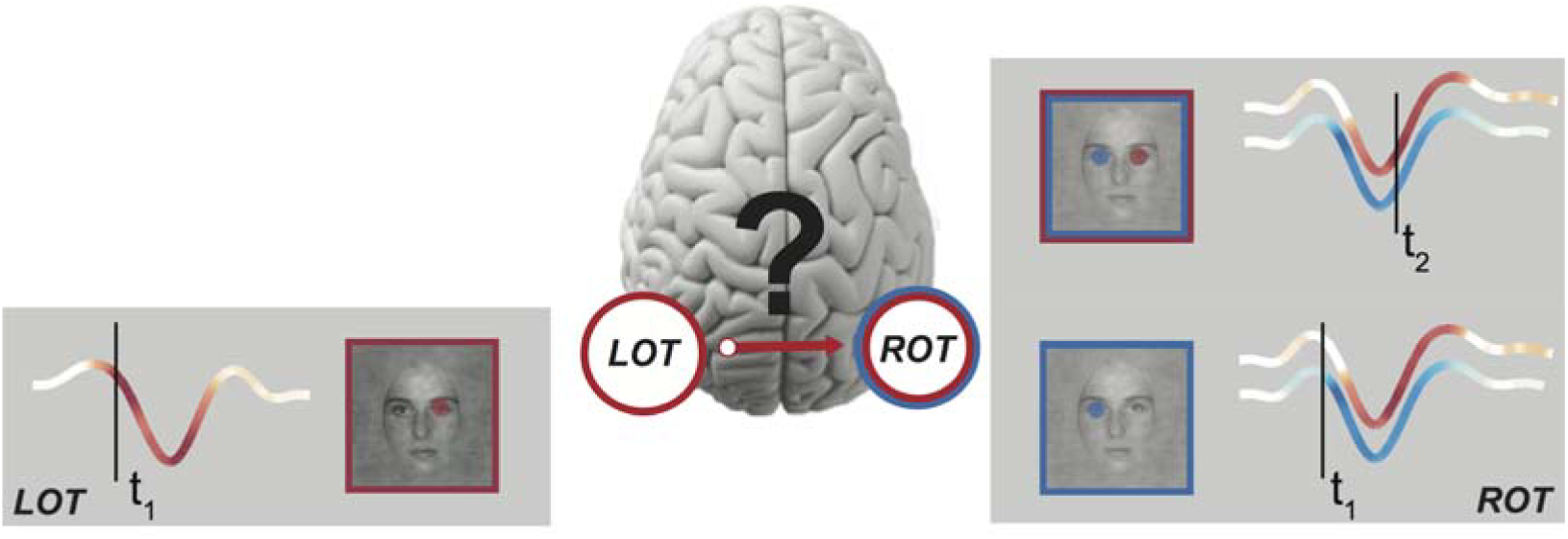
*Hypothesis of cross-hemisphere feature transfer along the N170 time-course.* At time t_1_, the Left Occipito Temporal (LOT) and the Right Occipito Temporal (ROT) sensors reflect coding of the contra-lateral left and right eye, respectively. Coding strength is represented with variations of hue (blue for the left eye; red for the right eye) directly on the LOT and ROT N170 ERP waveforms. At a later time t_2_, ROT also codes the ispi-lateral right eye. The anatomy of the visual system suggests that this sensitivity could arise from cross-hemisphere transfer from LOT, where the right eye is coded at t_1_.

Thus, converging evidence from face detection and categorization reveals that early face coding on the N170 differs between the left and right hemispheres. Specifically, as illustrated in Figure 1, at time t_1_ the right eye (represented in red) is initially coded on the left hemisphere N170, and the left eye (represented in blue) is coded on the right hemisphere N170 (Smith et al., 2007; Rousselet et al., 2014a). Furthermore, the later part of the N170 waveform additionally codes the eye ipsi-lateral to the sensor at time t_2_ (i.e. the right eye on the right sensor; the left eye on the left sensor, (Smith et al., 2007; Rousselet et al., 2014a)).

We know from anatomy and physiology that the visual system is lateralized across two hemispheres, with a separate visual hierarchy in each that processes the contra-lateral visual hemifield, from early to higher order visual areas, where processing becomes bi-lateral (Essen et al., 1982; Clarke and Miklossy, 1990; Saenz and Fine, 2010). Could the later N170 ipsi-lateral eye coding at t_2_ reflect the transfer of specific features coded at t_1_, across the hemispheres, through to high-level visual areas?

Figure 1 illustrates our hypothesis in the context of a face detection task, where the eyes (predominantly the left one) modulate reaction times (Rousselet et al., 2014a). Panels LOT and ROT (for Left and Right Occipito-Temporal sensors, respectively) illustrate initial coding of the contra-lateral eye at early time t_1_, closely followed by coding of the eye ipsi-lateral to the sensor at a later time t_2_—e.g. the right eye on ROT. Later coding of the right eye at t_2_ could arise from a causal transfer from its earlier coding at t_1_ on LOT on the opposite hemisphere. A demonstration of such feature transfer would suggest a reinterpretation of the N170, from a local event often interpreted as coding the entire face (Bentin et al., 1996; Eimer, 2000), to the reflection of a more global information processing network that spans both hemispheres across multiple stages of face coding in the visual hierarchy. Here, we demonstrate that the N170 does indeed reflect network-level information processing mechanisms.

In a face detection task (Rousselet et al., 2014a, 2014b), we instructed observers (N = 16) to detect on each trial the presence of a face sparsely and randomly sampled with small Gaussian apertures, see Figure 2A and (Rousselet et al., 2014a). Half of the trials sampled face images; the remaining half sampled amplitude spectrum matched noise, to dissociate spatial attention to feature location from feature coding per se. We recorded each observers’ EEG and face detection responses (correct vs. incorrect and Reaction Times, RTs (Rousselet et al., 2014a, 2014b)).

**Figure 2.**
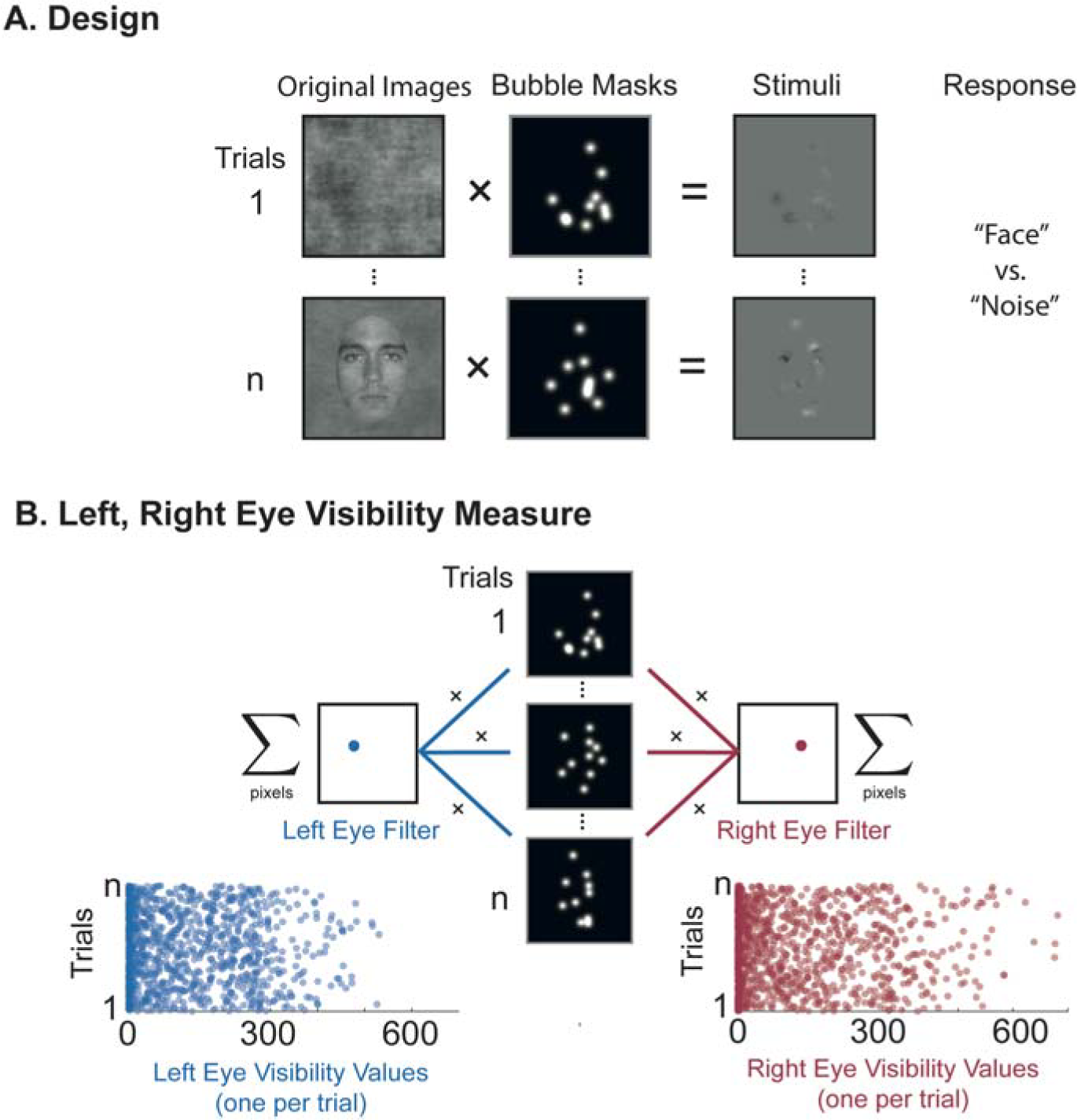
*Bubble sampling and eye visibility.* <u>A. Design.</u> On each trial, we used a bubble mask comprising 10 Gaussian apertures to sample visual information from either a texture or a face image. Observers pressed a key to indicate which they detected (“face” or “noise”). <u>B. Left, Right Eye Visibility Measure</u>. For each trial, we applied to the bubble mask a filter covering the spatial region of the left eye (left eye filter) and the right eye (right eye filter), counting the number of pixels the bubbles revealed within each regions. This produced two scalar values per trial representing the visibility of the left and right eye, respectively.

## Results

### Behavior

Observers were both fast and accurate, median of the median RT = 376 ms, [range = 287, 492]; mean accuracy = 91%, [range = 84, 97]. We used Mutual Information (MI) to compute the association between the sampled pixels and observer detection responses (face vs. noise). This revealed a significant relationship between pixel variations and behavior indicating that the pixels representing the left eye region in the image are relevant for behavior (Rousselet et al., 2014a Fig. 3). Computation of MI between sampled face pixels and the more sensitive RT measure revealed that most observers responded faster on trials that revealed the left eye—a minority also responded faster to trials revealing the right eye (Rousselet et al., 2014a Fig. 3). On noise trials, MI values were low and not clustered on specific face features. Henceforth, we focus on the EEG coding and transfer of the eyes on face trials due to their prominence across observers in the face detection task (Rousselet et al., 2014a).

**Figure 3.**
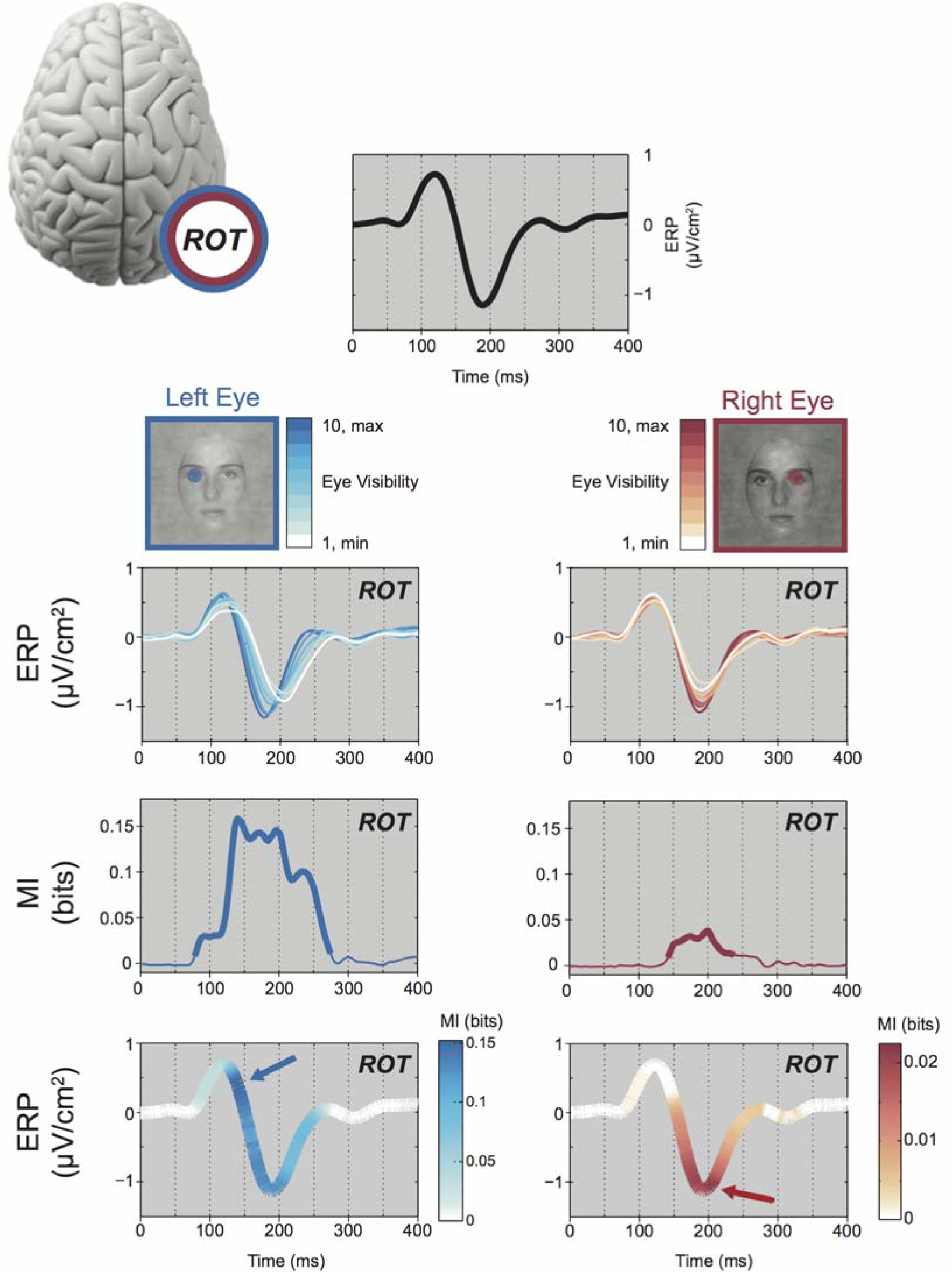
*Coding.* The top panel illustrates the ERP measured on ROT for a typical observer (black curve). For this observer, ROT was selected as electrode B9 in the Biosemi coordinate system, which is posterior to PO8 (on the same radial axis). The left eye and right eye schematics illustrate with their color-coded blue and red scales the deciles of eye visibility across experimental trials (with 10 = highest eye visibility trials and 1 = lowest). Directly below, we recomputed ROT ERPs using only the trials from each decile of left and right eye visibility, to illustrate how changes of eye visibility modulates ERPs. The next row of curves show the MI between left eye (blue curve) and right eye (red curve) visibility and corresponding ROT EEG response (a thicker line indicates regions of statistically significant MI, referred to as left and right eye coding). Finally, the bottom panels illustrate the MI curves reported on the ERP curve to precisely indicate when, over the time course of the ERP, eye coding peaks (indicated with a color-coded arrow).

### EEG

We removed one observer from analysis due to poor EEG signal. For the remaining 15 observers and for each sensor we computed the sensitivity of the EEG to the left and right eyes as follows. First, we used a mask for the left and the right eye regions and summed the Gaussian apertures within each eye region. The resulting two scalar values represent the visibility of each eye on each trial (Figure 2B). Then, across trials we quantified the relationship (MI) between eye visibility values and the Current Source Density EEG measured at each sensor and time point—henceforth we refer to these interchangeably as MI time courses or eye coding curves. To remove any effects of weak statistical dependence between the left and right eye sampling, throughout when we refer to MI about an eye feature we actually calculated Conditional Mutual Information (CMI), conditioning out any effect of the visibility of the alternate eye. On each hemisphere, we then identified the single occipital-temporal sensor (i.e. LOT and ROT) with largest MI to the contralateral eye within the N170 time window (100-200ms)—see Materials and Methods.

To address our hypothesis (cf. Figure 1), we propose three requirements that are necessary for the existence of a causal transfer of stimulus features between two brain regions. The first requirement is *coding:* Both regions should code the same stimulus feature (e.g. the right eye at t1 on the LOT N170; the right eye at t_2_ on the ROT N170). The second requirement is *temporal precedence:* Feature coding in the first region should occur before coding of the same feature in the other region (e.g. the right eye on LOT N170 at t_1_ and the right eye on ROT N170 at t_2_). Finally there must be *coding equivalence:* Feature coding should be the same in both regions (e.g. coding of the right eye at t_1_ on LOT and at t_2_ on ROT should be the same).

### Coding

To operationalize coding, we refer to the MI time courses of the chosen LOT and ROT sensors for the left and right eye. Figure 3 illustrates this analysis for one observer. For reference, the black curve shows a typical N170 obtained on the ROT sensor. Standard interpretations would consider this average as a local response to full-face stimuli, in contrast to other categories (Bentin et al., 1996; Eimer, 2000). Here, random sampling with bubbles changes the visibility of the left and right eyes across trials (Figure 2B) and so we can analyze how eye visibility modulates singletrial N170 responses. On the ROT N170, we split the trials into 10 bins of left eye visibility (deciles of the distribution across trials; represented with shades of blue) and right eye visibility (represented with shades of red). For each bin we computed and plotted the mean ERP. Figure 3 illustrates that increased left eye visibility causes an earlier and larger N170. Increased visibility of the right eye caused larger N170 amplitude, with no change in latency.

Plotting the ERP for bin of eye visibility demonstrates a modulation. To quantify this coding, we calculated, for each eye, the MI between eye visibility and the corresponding EEG response at each time point at LOT and ROT sensors (see Methods). In Figure 3, MI time courses indicate with a thicker line the time windows of a statistically significant relationship (p=0.01, corrected for multiple comparison over time [0-400ms], all sensors and the two eye features with the method of maximum statistics). MI time courses show that in this observer the ROT N170 codes the left and right eyes. All 15 observers showed significant MI on the contra-lateral sensor for at least one eye (13/15 significant for both eyes). 14/15 observers also showed significant MI of the eye ipsi-lateral to the sensor, for at least one eye (13/15 significant for both eyes). These results satisfy the coding requirement because across observers and stimulus features we found 26 instances (out of 30 = 15 subjects x left/right eye) of significant eye MI on both contra-lateral and ipsi-lateral sensors.

### Temporal Precedence

In Figure 3, the lower plots report the eye coding MI curves superimposed as hue on the ROT ERP time-course, to directly visualize the timing of left and right eye coding on this sensor. The ROT N170 codes the contra-lateral left eye throughout its time, with a strongest early effect at the onset of the negative deflection (see blue arrow). The ROT N170 also codes the ipsi-lateral right eye, but only later, with the strongest effect just after the N170 peak (see red arrow). This illustrates that the ROT N170 codes the contra-lateral left eye before the ipsi-lateral right eye.

We now demonstrate the temporal precedence of the left (or right) eye across the contra and ipsi-lateral sensors. To visualize this comparison, Figure 4A reveals the peak-normalized MI time courses of the left eye on ROT (plain blue line) and LOT (dashed blue line); Figure 4B presents the MI of the right eye on LOT (plain red line) and ROT (dashed red line). Comparison of the solid (contra-lateral sensor) to the dashed (ipsi-lateral sensor) MI curves illustrates contral-lateral temporal precedence in both cases. To quantify temporal precedence, we considered each instance (specific observer and eye feature) with significant same eye coding across the contra‐ and ipsi-lateral sensors (N = 26). For each instance, we quantified the coding latency by normalizing the MI curves to their peak values and calculating the average delay between the MI curves over the y-axis region where both were significant (grey region, Figure 4C; see Materials and Methods). At the group level, observers showed a significant temporal precedence of same eye coding on the contra-lateral before the ipsi-lateral sensor (Figure 4D; Sign Rank test, p=7.4e^-7^). The median contra-to-ipsi coding delay was 15.7 ms (1^st^ quartile = 7.7, 3^rd^ quartile = 25.5 ms). 13/15 observers have significant bilateral coding with a contra-ipsi latency of greater than 7 ms for at least one eye feature. Across observers, we have now established temporal precedence of same eye coding from the contra-lateral to the ipsi-lateral sensor.

**Figure 4.**
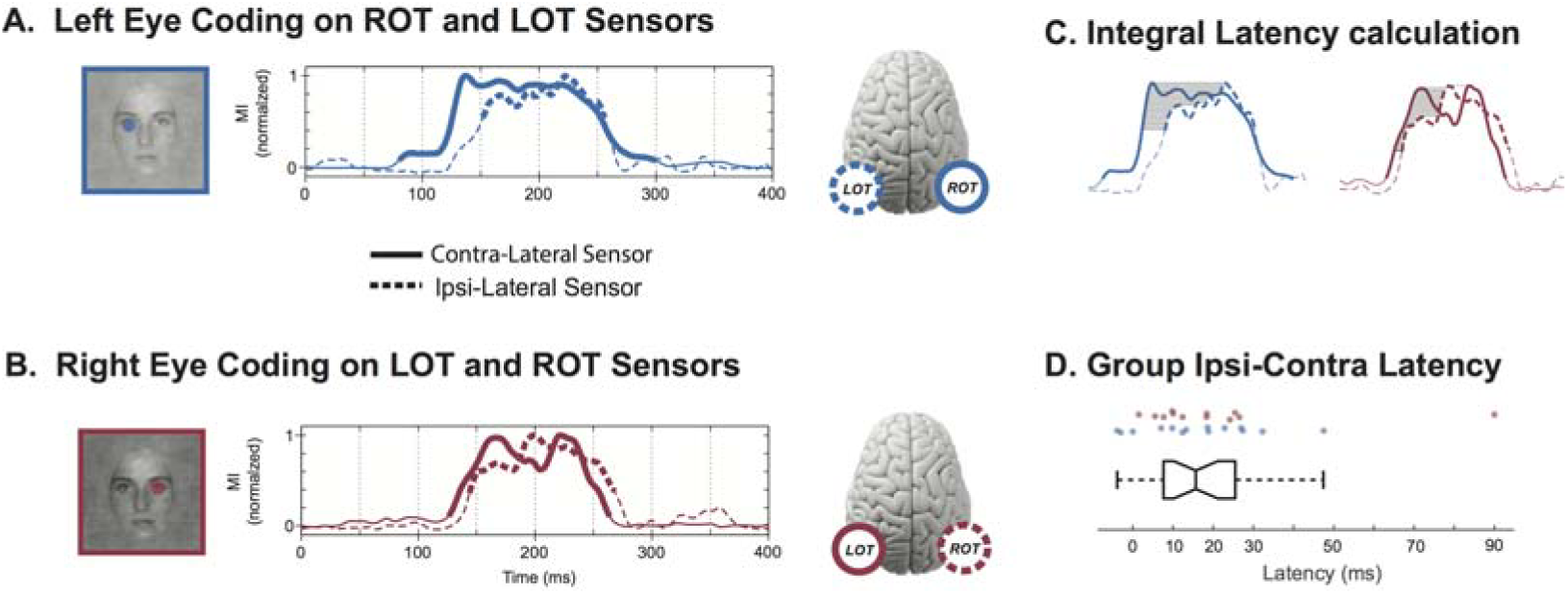
*Latency.* <u>A. Left Eye Coding on ROT and LOT Sensors.</u> The peak-normalized MI curves indicate the initial coding of the left eye on the contra-lateral ROT sensor (plain curve) and the later coding of the same eye on the ipsi-lateral LOT sensor (dashed curve). Thicker sections of the MI curves represent statistical significance. <u>B. Right Eye Coding on LOT and ROT Sensors.</u> The normalized MI curves indicate the initial coding of the right eye on the LOT sensor (plain curve) and the later coding of the same eye on the ROT sensor (dashed curve). Thicker sections of the MI curves represent statistical significance. <u>C. Integral Latency Calculation (see Methods).</u> Illustration of the eye coding latency calculation (left eye, blue; right eye, red) from the normalized MI curves of the contra‐ and ipsi-lateral sensors (solid, dashed lines respectively). <u>D. Group Ipsi-Contra Latency.</u> Box plot of the group latency (in ms) between contra‐ and ispi-lateral coding of the same eye across hemispheric sensors. Positive values correspond to earlier contra-lateral coding. Each dot above represents a particular observer and eye feature—blue for left eye (ROT to LOT latency), red for right eye (LOT to ROT latency).

### Coding Equivalence

We turn to coding equivalence, the third and final necessary condition for causal feature transfer. Figure 5A schematizes our results so far. We know that the LOT N170 codes the contra-lateral right eye at t_1_—this is represented with plain lines. We also know that the ROT N170 codes the same eye at a later time, t_2_—this is represented with dashed lines. Now, we establish that the eye feature coded at these two different time points and on different sensors is mostly equivalent.

**Figure 5.**
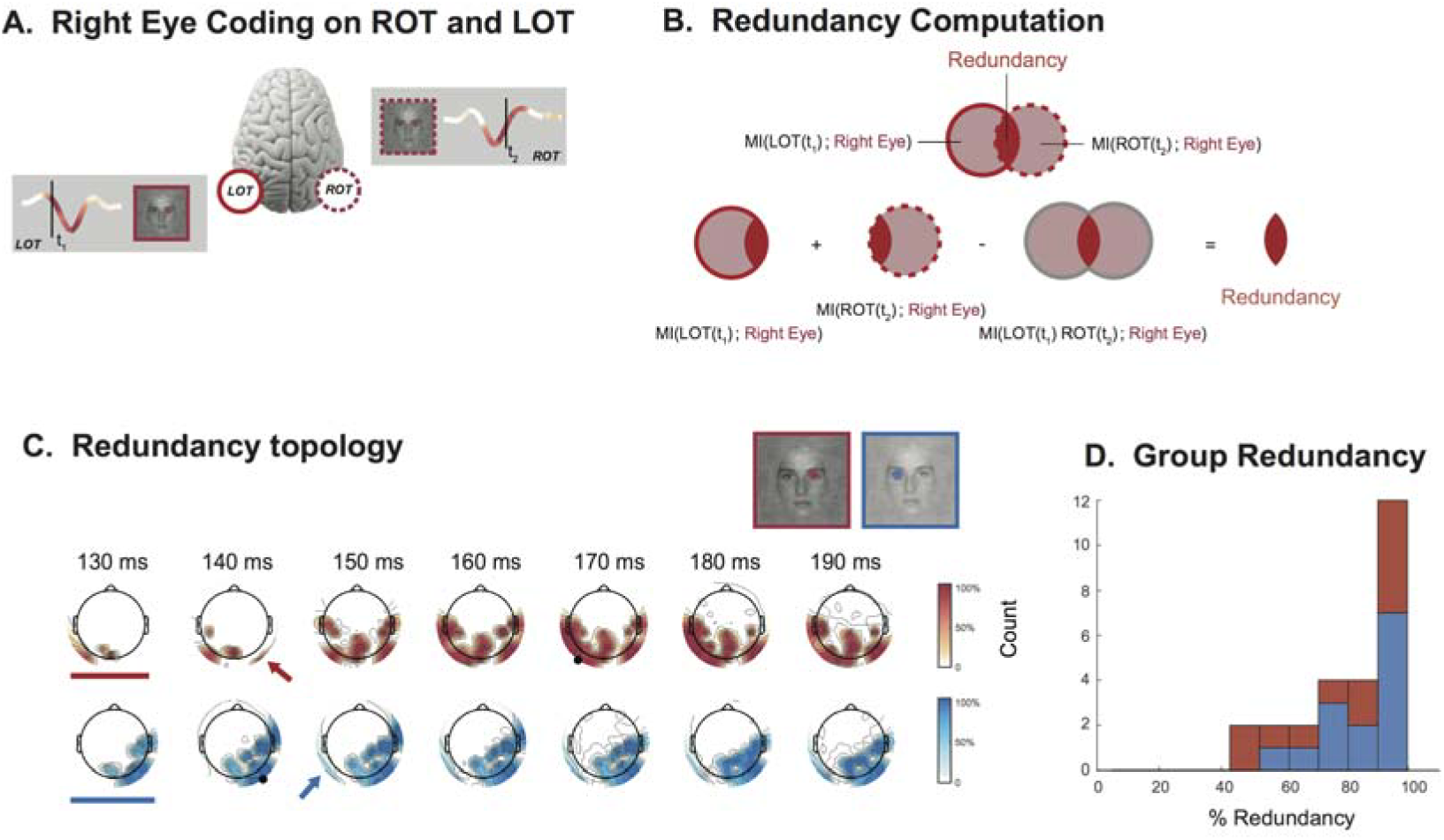
*Redundancy.* <u>A. Illustration of the coding redundancy of the right eye between early LOT and late ROT.</u> Coding redundancy of the right eye is computed between LOT and ROT at the peak of each MI curve (i.e. at t_1_ on LOT and t_2_ on ROT). <u>B. Redundancy Computation.</u> Venn diagrams illustrate the computation of redundant (i.e. intersecting) eye information on LOT and ROT. <u>C. Redundancy Topography.</u> Using as seed the peak MI time point on LOT (marked with black circle on red topographies), and ROT (black circle on blue topographies) the time courses of redundancy topographies illustrate when and where (see color-coded arrow) redundant ipsi-lateral coding of the eyes begins. Note that for both eyes, the 130ms time point (indicated with bar) shows no ipsi-lateral redundancy. <u>D. Group Redundancy.</u> Left (in blue) and right (in red) eye coding redundancy between LOT and ROT, at the respective peaks of the MI curves, expressed as percentages.

To provide an intuitive understanding of coding equivalence, consider Figure 5A and a putative observer who would read out only the early EEG responses from the LOT sensor. Could they predict the visibility of the right eye on each trial?

Would adding information from the later EEG responses on ROT improve their prediction? If the early LOT and later ROT information were the same, adding the later ROT responses would not add any new knowledge about right eye visibility and thus would not improve prediction. However, if adding ROT information did improve prediction, then ROT responses would contain extra information about the visibility of the eye that is not already available in LOT responses, indicating that ROT and LOT coding of the right eye were not equivalent.

We formalized such coding equivalence with an information theoretic quantity called *redundancy* illustrated in Figure 5B (see Methods). Venn diagrams represent the MI (at peak time) to the right eye on LOT (plain line) and ROT (dashed line). Coding equivalence corresponds to their overlap (i.e. their intersection). Because MI is additive we measure this overlap directly, with a simple linear combination of MI quantities. We sum the LOT and ROT MI separately (which counts the overlapping MI twice) and subtract the MI obtained when LOT and ROT responses are considered together (which counts the overlap once). The resulting redundancy value measures coding equivalence—i.e. the MI that is shared between the LOT and ROT responses.

For each observer, we computed redundancy between the N170 MI peaks for for the left eye (a transfer from right to left hemispheres) and for the right eye (the opposite transfer, from left to right hemispheres). Figure 5C illustrates the redundancy topography time courses for one typical observer. Redundancy is computed with respect to a seed response—a particular time point (MI peak) of either LOT (for right eye redundancy, red topographies, seed indicated with black marker) or ROT (for left eye redundancy, blue topographies, seed indicated with black marker). At 130ms there is contra-lateral, but no ipsi-lateral coding of either eye (indicated with bars); ipsi-lateral redundant coding appears 10-20ms later (arrows). The redundancy topographies also illustrate that our analyses do not depend on the choice of a single seed sensor in each hemisphere. Figure 5D shows a histogram of the normalised redundancy over all 15 observers. In almost all cases redundancy is large (median = 88%) demonstrating the high coding similarity of the same eye across hemispheric locations and time segments of the N170. 14/15 observers have contra-ipsi redundancy greater than 50% for at least one eye feature (21/26 bi-lateral instances).

To provide further evidence that delayed redundancy across hemispheres actually represents feature transfer between the two regions, we calculated the MI between LOT and ROT peaks directly, conditioning out the effect of variability of both eye features. These values were large (compared to stimulus feature coding) and significant. All 26 instances showed significant conditional MI: median = 0.32 bits, min = 0.07 bits, max = 0.65 bits; median of the lower boundaries of the 99% bootstrap confidence interval = 0.25 bits, min = 0.04 bits, max = 0.55 bits). This demonstrates that there is a time-delayed relationship between LOT and ROT on single trials, over and above the relationship that could be expected if these two regions received a common eye feature signal from a third region but did not exchange information with each other. This can be interpreted as causal transfer within the Wiener-Granger framework (Wiener, 1956; Granger, 1969; Bressler and Seth, 2011), and so shows that our observations are better explained by a model that includes genuine transfer of eye information between the regions, rather than a model involving a third region that sends the same eye signal with a different delay to LOT and ROT.

## Discussion

In a face detection task, we derived an abstract, high-level information processing interpretation of brain activity. First, using the Bubbles procedure coupled with behavioral responses, we showed *what* information supports face detection: the eyes of a face. Then, with the Bubbles procedure coupled with CSD transformed EEG data to improve spatial localization (Tenke and Kayser, 2012), we reconstructed a network revealing that the sources generating the N170 code and transfer the eyes of a face. We specified three necessary conditions for inferring feature transfer within our measurement paradigm: *coding* of the same features in two brain regions, *temporal precedence* of coding in one region with respect to the other and *redundancy* of feature coding between the two regions. Our analyses revealed that 13/15 observers individually meet all three conditions. Specifically, in each observer we demonstrated *coding* of the same eye on the LOT and ROT sensors. We showed the *temporal precedence* of same eye coding at contra-before ipsi-lateral sensors. Finally, we showed high *redundancy* (i.e. high coding similarity) between early and late coding of the same eye on opposite hemispheres. Together, these three conditions suggest causal transfer of eye information in the Wiener-Granger sense (e.g. right eye from the early part of the LOT N170 to late part of the ROT N170). Though we selected one LOT and one ROT electrode per observer for statistical analyses, the topographic maps of Figures S2-S16 reveal that the three conditions would be met by choosing other electrodes from the extended lateral regions of delayed redundancy across hemispheres. The clear separation between the clusters of redundant information on the left and right hemispheres further supports the claim that LOT and ROT signals originate from different brain regions.

The N170 now appears as a deceptively simple signal that actually reflects an inter-hemispheric information processing network that codes and transfers face features. A distinctive feature of our approach is that we are not just reconstructing a network on the basis that two brain regions communicate; we are revealing *what they are communicating about—i.e.* the information content underlying face detection. It is the novel quantification of the content of information coding and transfer that represents an important step towards a new brain algorithmics to model the information processing mechanisms of perception and cognition (Schyns et al., 2009). This is a radical departure from typical N170 quantifications and interpretations, which compare peak amplitude and latency differences between the average EEG responses to a few categories comprising complete stimulus images (Bentin et al., 1996; Rossion and Jacques, 2008). In contrast, here we decompose the stimulus with Bubbles to test the N170 responses against random samples of stimulus pixels presented on each trial. This enables a precise characterization of the information subtending the task and the dynamics of coding alongside the N170 time course in individual observers. We demonstrated that focusing on the average N170 peak was restricting interpretation of the N170 signals because (a) feature coding is represented in the single trial variance of the N170 (not its mean), revealing that (b) contra-lateral eye coding starts on the N170 downward slope, ~40 ms prior its peak (see also (Schyns et al., 2007)) and that (c) the rebound from the peak codes the transferred ipsi-lateral eye from the other hemisphere. Together, our results demonstrate that the N170 cannot be fully interpreted as an isolated event on one hemisphere. Rather, it reflects the coding and transfer functions of an information processing network involving both hemispheres.

### Feature Transfer

Our results provide strong evidence for causal feature communication between hemispheres. We review the evidence in turn. First, the lateralized anatomy of the visual system prescribes that the first cortical processing of each eye should occur in the early part of the contra-lateral visual hierarchy. This is congruent with our findings. As there is no direct anatomical pathway from the ipsi-lateral hemifield (only a contra-lateral pathway), the ipsi-lateral eye information present in the second part of the LOT and ROT N170 should be transferred from the opposite hemisphere. We demonstrated this functionally, without recourse to anatomical priors in our analysis. Second, the latency timings of inter-hemispheric feature transfer (median 11 ms) are consistent with previously reported inter-hemispheric transfer times (Brown et al., 1994; Ipata et al., 1997). Third, we demonstrate that the coding redundancy between LOT and ROT occurs in the context of single-trial temporal relationships between these regions that cannot be explained simply by a third region sending a common eye feature signal with a different delay. In sum, the reported evidence is consistent with a transfer between the two hemispheres, a proportion of which is *about* the eye.

### Sources Generating the N170

Further research will seek to reduce the abstract information dynamics presented here. We will examine the networks of brain sources that generate the left and right hemisphere N170s and implement the information processing functions of task-dependent contra-lateral and ipsi-lateral feature coding and cross-hemispheric transfer. In our data, topographic maps of contralateral eye coding suggest the involvement of posterior-lateral sources. Source analyses or correlations between BOLD and ERP amplitudes suggest that the N170 sources locate around the STS (Watanabe et al., 2003; Itier and Taylor, 2004; Sato et al., 2008; Nguyen and Cunnington, 2014), the fusiform gyrus (Horovitz et al., 2004), or both (Sadeh et al., 2010; Dalrymple et al., 2011; Prieto et al., 2011). To better localize these sources and understand how they code and transfer task-relevant features, we could apply our bubbles paradigm with single-trial fMRI-EEG measures. In fact, an MEG reverse-correlation study revealed coding of face features, including the eyes, in the time window of the M170 in lateral cortical areas (Smith et al., 2009). Finally, intracranial data also support the involvement of occipital and temporal lateral areas, such as the right inferior occipital gyrus, to generate scalp N1/ N170 (Sehatpour et al., 2008; Rosburg et al., 2010; Engell and McCarthy, 2011; Jonas et al., 2012, 2014). So, lateral sources are likely to be involved in the generation of the N170. The information processing mechanisms revealed here guide their study at the source level, in terms of the timing of coding and transfer of specific features.

### Advantages of Information Theory for Analysis of Brain Signals

It is often thought that the EEG signal is too noisy for single-trial analyses as done here. It is worth noting that several novel methodological developments delivered important advances. First, a copula-based MI estimator provided a flexible and powerful multivariate statistical framework to quantify feature coding over the full temporal resolution of EEG (Ince et al., 2015). Second, as an improved measure of EEG activity, we considered the bivariate variable consisting of the recorded voltage at each time point together with the temporal gradient (see Methods and Figure S1). Third, to eliminate effects of the alternative eye when computing eye coding, we used conditional MI throughout our analyses. Conditional MI is a powerful and rigorous approach to quantify feature coding when stimulus features are correlated, and is applicable in any sensory modality. Finally, the additivity of MI enables a direct quantification of feature coding interactions between different regions of the brain and time points of their activity. Here, we used MI additivity to compute redundancy, but the same computation can also reveal synergistic interactions. Using MI to quantify coding redundancy and synergy has broad applications in neuroimaging, from quantifying the interactions between different spatial regions and time points to construct brain representations (as considered here), to quantifying the relationships between signals from different brain imaging modalities (e.g. EEG/fMRI).

### Hierarchical Analysis of Information Processing

Our interpretative approach can be applied hierarchically, to different levels of granularity of response (e.g. from behavior to EEG to neurons), to quantify information processing mechanisms at each level. At the coarsest level, with behavioral measures (accuracy and RT) we determine *what* visual features the organism processes to discriminate faces from noise—i.e. the two eyes. This reduces the full high-dimensional stimulus to a few, lower-dimensional task-relevant features. Going down the hierarchy, with integrated EEG measures we determine *where, when* and *how* states of brain activity (e.g. variance of the early and late parts of the single-trial N170 signals) code and transfer task-relevant features. That is, we relate information processing to a few states of brain activity (e.g. coding of the contra-lateral eye during the early part of the ROT and LOT N170; coding of the ipsi-lateral eye over the later part). We also relate operations on information (e.g. transfer of the ipsi-lateral eye across hemispheres) to brain state transitions (e.g. eye transfer occurs between the early and late part of the LOT and ROT N170s). Experimentally, we can repeat the exercise one (or several) hierarchical level(s) down, for instance measuring the localized MEG sources that generate the N170, to add the details of information processing that would be reverse engineered from the finer grained measures of neural activity. We expect these more detailed information processing descriptions to be consistent with the results described here.

The critical point is that while the information processing ontology produces finer information processing details with increasing granularity of brain measures, it preserves the information gained from analyses at more abstract levels (as shown here between behavior and the EEG that preserves the eyes). Abstract levels guide the search for detailed implementation of the behaviorally relevant information (e.g. contra‐ and ipsi-later eyes) and functions (e.g. coding, transfer) in the increased complexity of the lower levels—i.e. whatever else the neural populations in the sources of the N170 may be encoding, the population must be sensitive to first the contra‐ and then the ipsi-lateral eye and transfer of the latter should come from a population on the other hemisphere. Our approach is similar to analyzing a computing architecture hierarchically across levels, from the most abstract (e.g. “send mail”), to its programming language algorithm, to its assembly language, to its machine-specific implementation. The critical difference between a brain and a computer is that whereas we can directly engineer (via a hierarchy of compilers) an abstract information algorithm into a computer hardware, we can only reverse engineer an information processing algorithm from brain data to infer a hierarchy.

### Influences of Categorization Tasks

Our approach will be particularly interesting when applied to study the processing of the same stimuli when the observer is performing different tasks. For example, with multiple categorizations of the same faces (e.g. gender, expression and identity), we could determine from behavior the specific features that are relevant for each task (the *what*) and then trace, as we have done here, *where* and *when* each feature set is coded and transferred between localized brain sources (e.g. with MEG (Smith et al., 2009)). As the task and the associated behaviorally relevant information changes, we can determine how the corresponding processing in brain networks is affected: Where and when are task-relevant features coded and task-irrelevant features suppressed? This is a pre-requisite to addressing elusive questions such as the locus of selective attention, the role of top-down priors, and their influence on the construction of the information content of stimulus perception and categorization. How does network feature communication change with task? Again, this is an important condition to understand for example information integration. Our results propose a specific timing on feature coding and transfer that could constrain the study of feature integration mechanisms—specifically, these should occur after transfer of the contra-laterally coded features, possibly in occipito-temporal sources. Can we relate selective coding of behaviorally relevant features to specific decision-making processes (Smith et al., 2004; Philiastides and Sajda, 2006)(Philiastides and Sajda, 2006)?

Here, by addressing the what, where, when and how questions of information processing, we proposed a radically new interpretation of the N170, as reflecting a network that codes and transfers a specific information content across hemispheres. The main methodological advantage of focusing on the *what* and then reducing its processing across the levels of an information processing ontology is akin to the main recommendation of Marr’s computational analysis: The abstract information goals of the system guide the analysis. Revealing the information, from behavior to the processing states of the brain and their transitions brings us one step closer to the ultimate goal of cognitive neuroimaging of understanding the brain as a machine that processes information.

## Materials and Methods

The data considered here were already reported in (Rousselet et al., 2014a). Full experimental details are provided there. Data are available at (Rousselet et al., 2014b).

### Observers

The study comprised 16 observers: 9 females, 15 right-handed, median age 23 (min 20, max 36). Prior to the experiment, all observers read a study information sheet and signed an informed consent form. The experiment was approved by the Glasgow University College of Science and Engineering Ethics Committee with approval no. CSE00740. All observers had normal or corrected-to-normal vision and contrast sensitivity of 1.95 and above (normal score).

### Stimuli

Stimuli were gray-scale pictures of faces and textures (Rousselet et al., 2014a Fig. 1). Faces from 10 identities were used; a unique image was presented on each trial by introducing noise (70% phase coherence) into the face images (Rousselet et al., 2008). Textures were face images with random phase (0% phase coherence). All stimuli had an amplitude spectrum set to the mean amplitude of all faces. All stimuli also had the same mean pixel intensity, 0.2 contrast variance and spanned 9.38 x 9.38 degrees of visual angle. The face oval was 4.98 x 7.08 degrees of visual angle. Face and noise pictures were revealed through 10 two-dimensional Gaussian apertures (sigma = 0.368 degrees) randomly positioned with the constraint that the center of each aperture remained in the face oval and was at a unique position. In the rest of this article, we refer to these masks with Gaussian apertures as *bubble masks*.

### Experimental Procedure

At the beginning of each of two experimental sessions, we fitted observers with a Biosemi head cap comprising 128 EEG electrodes. We instructed observers as to the task, including a request to minimize blinking and movements. We asked observers to detect images of faces and textures as fast and as accurately as possible. They pressed the numerical pad of a keyboard for response (‘1’ for face vs. ‘2’ for texture) using the index and middle fingers of their dominant hand. Each experimental session comprised 1,200 trials, presented in blocks of 100, including 100 practice trials. All observers participated in two experimental sessions lasting in total about 4 hours and bringing the total number of trials per observer to 2,200.

Each trial began with the presentation of a small black fixation cross (0.48 x 0.48 degrees of visual angle) displayed at the center of the monitor screen for a random time interval between 500 to 1000 ms, followed by a face or texture image presented for ~82 ms (seven refresh frames). A blank gray screen followed stimulus presentation until observer response.

### EEG Preprocessing

EEG data were re-referenced offline to an average reference, band-pass filtered between 1Hz and 30Hz using a fourth order Butterworth filter, down-sampled to 500 Hz sampling rate and baseline corrected using the average activity between 300ms pre-stimulus and stimulus presentation. Noisy electrodes and trials were detected by visual inspection on an observer-by-observer basis. We performed ICA to reduce blink and eye-movement artifacts, as implemented in the infomax algorithm from EEGLAB. Components representing blinks and eye movements were identified by visual inspection of their topographies, time-courses and amplitude spectra. After rejection of artefactual components (median = 4; min = 1; max = 10), we again performed baseline correction. Finally, we computed single-trial spherical spline Current Source Density (CSD) waveforms using the CSD toolbox with parameters iterations = 50, m = 4, lambda = 1.0e-5 (Kayser and Tenke, 2006; Tenke and Kayser, 2012). The CSD transformation is a 2^nd^ spatial derivative (Laplacian) of the EEG voltage over the scalp that sharpens ERP topographies and reduces the influence of volume-conducted activity. The head radius was set to 10 cm, so that the ERP units in all figures are μV/cm^2^. We also calculated the central-difference numerical temporal derivative of the CSD signal for each sensor and on each trial.

### Behavior Information: Mutual Information between Pixels and Detection and Reaction Time

Our analysis focuses on the single trial bubble masks because they control the visibility of the underlying image. Bubble masks take values between 0 and 1, controlling the relative opacity of the mask at each pixel, with 0 being completely opaque (pixel was shown grey) and 1 being completely translucent (pixel of underlying image was shown unaltered). We analyzed the bubble masks at a resolution of 192 x 134 pixels. For each observer and for each image pixel, we applied Mutual Information (MI) (Cover and Thomas, 1991) to compute the relationship between single-trial pixel visibility and detection responses, and separately the relationship between single-trial pixel visibility and reaction times. We computed one MI pixel image for the face trials and a separate MI image for the texture trials. We found (Rousselet et al., 2014a Fig. 3) that the main face regions that systematically produced an effect on behavior were the eyes of the face images, but there was no such effect on texture trials. In the ensuing EEG analyses, we therefore only considered the face trials.

### EEG Information: Mutual Information between Eye Visibility and EEG Sensor Response

As the eyes are the critical regions affecting behavioral measures, we reduced the dimensionality of the single trial bubble masks to a measure of the visibility of each eye, to directly track their coding in the EEG. To this aim, we manually specified two circular spatial filters, one to cover each eye region (determined from the mean face image). We applied each of these filters to the bubble mask on each trial to derive a scalar value representing the visibility of the left eye and another scalar value representing the visibility of the right eye. Specifically, for each trial we sum the total bubble mask visibility within the left eye region (and separately for the right eye region). These two scalars represent the area of each eye that is was visible on that trial (Figure 2B). We used the same eye region filters for all observers.

For each observer, we then used MI to quantify the relationship between the scalar visibility of each eye on each trial and the corresponding EEG signal, on each electrode (median 118, range 98-125). MI is a statistical quantity that measures the strength of the dependence (linear or not) between two random variables. It can also be viewed as the effect size for a statistical test of independence. One advantage of MI is that it can be applied with multivariate response variables, as done here. For each sensor and time point we considered a bivariate EEG response comprising the raw voltage value and the instantaneous gradient (the temporal derivative of the raw EEG). Supplemental Figure S1 illustrates with an example the smoothing effect of including the instantaneous gradient in the MI calculation. The top plot reproduces the panel of Figure 3 showing how visibility of the left eye modulates the ERP waveform. The rank correlation plot illustrates rank correlations between left eye visibility and EEG voltage measured at each time point. Correlation is a signed quantity that reveals transitions between regions of positive correlations (when ERP voltage increases with increased eye visibility) and regions of negative correlations (when ERP voltage decreases with increased eye visibility). At each transition (zero crossing), there is no measurable effect in the raw EEG voltage, even though this time point is within the time window when eye visibility clearly affects the ERP waveform. With MI, we addressed this shortcoming by computing at each time point the relationship between eye visibility and the bivariate EEG response comprising the raw EEG voltage and its instantaneous gradient. The two bottom MI plots demonstrate that the effect of adding the instantaneous gradient to smooth out the transitions and provide a more faithful and interpretable measure of coding dynamics.

The sampling strategy using bubbles (with fixed number of 10 bubble apertures per bubble mask on each trial) induces a weak dependence between the two (left and right) eye feature values across trials (median MI = 0.011 bits, range 0.0015-0.023 bits, significant at p = 0.01 uncorrected for 12/15 observers). This arises because a high visibility value of the left eye on a given trial indicates a high concentration of individual bubbles in that area, implying that fewer bubbles are distributed over the remainder of the face, including the right eye. This dependence introduces an ambiguity for interpretation. For example, if the EEG signal had a high left eye MI value and a low right eye MI value at a given time point, two interpretations would be possible. First, the EEG genuinely codes the right eye, irrespective of the visibility of the left eye. Second, the EEG only codes the left eye, and the low right eye MI value arises from the statistical dependence between the two eyes, as just discussed. We addressed this potential ambiguity with Conditional Mutual Information (CMI, the information theoretic analogue of partial correlation) (Cover and Thomas, 1991; Ince et al., 2012). CMI quantifies the relationship between any two variables (e.g. left eye visibility and the EEG response) while removing the common effect of a third variable (e.g. right eye visibility). We thus calculate CMI between left eye and the EEG signal, conditioned on the right eye (i.e. removing its effect), and similarly the CMI between the right eye and the EEG signal conditioned on the left eye (removing its effect). Throughout the paper whenever we refer to MI about eye visibility, we actually calculated CMI conditioned on the alternative eye.

We calculated CMI using a bin-less rank based approach based on copulas. Due to its robustness this approach is particularly well suited for noisy continuous valued neuroimaging data such as EEG and provides greater statistical power than MI estimates based on binning. The following paragraphs detail this MI estimator. They can be skipped without loss of continuity.

A copula (Nelsen, 2007) is a statistical structure that expresses the relationship between two random variables (e.g. between the left eye visibility and the EEG response on one electrode, at one time point). The negative entropy of a copula between two variables is equal to their MI (Ma and Sun, 2011). On this basis, we fit a Gaussian copula to the empirical copula obtained from the eye visibility and EEG response, estimate its entropy and then obtain MI as the negative of this entropy. While this use of a Gaussian copula does impose a parametric assumption on the form of the interaction between the two variables, it does not impose any assumptions on the marginal distributions. This is important because the distribution of the visibility of the eye across trials is highly non-Gaussian. Since the Gaussian distribution is the maximum entropy distribution for a given mean and covariance, the Gaussian copula has higher entropy than any other parametric copula model that preserves those statistics. This MI estimation is therefore a lower bound on the true MI.

In practice, we calculated the empirical CDF values for a particular sensor and time point by ranking the data recorded across trials, and then scaling the ranks between 0 and 1. We then obtained the corresponding standardized value from the inverse CDF of a standard normal distribution. We performed this normalization separately for the EEG voltage and gradient, before concatenating them to form a 2D EEG response variable *R*. We computed CMI between these standardized variables using the analytic expressions for the entropy of uni-, bi-, tri-and quad-variate Gaussian variables (Misra et al., 2005; Magri et al., 2009):

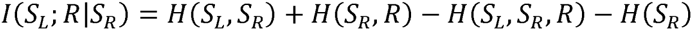

We estimated entropy terms and corrected for the bias due to limited sampling using the analytic expressions for Gaussian variables (Misra et al., 2005; Magri et al., 2009). A particular advantage of this estimation method is its multivariate performance, which we exploit here with our 2D EEG voltage and gradient responses.

We determined statistical significance with a permutation approach, and addressed the problem of multiple comparisons using the method of maximum statistics (Holmes et al., 1996). For each of 200 permutations, we randomly shuffled the recorded EEG data across trials and repeated the MI calculation for each sensor and time point. We computed the maximum of the resulting 3D MI matrix (time vs. EEG sensors vs. left and right eye visibility) for each permutation. For each observer we used the 99^th^ percentile across permutations as the statistical threshold for each observer.

### Selection of LOT and ROT Sensors

As described above we calculated MI between EEG and visibility of each eye for each sensor and time point. We selected for further analyses the lateral occipitotemporal sensors with maximum MI for the eye contra-lateral to the sensor in a time window 100 and 200 ms post stimulus. On the left hemisphere, for LOT we selected the sensor with maximum right eye MI from sensors on the radial axes of P07, P7 and TP7 (excluding midline Oz and neighboring O1 radial axes). On the right hemisphere, for ROT we selected the sensor with maximum left eye MI from sensors on the radial axes of PO8, P8, TP8 (excluding midline Oz and neighboring O2 radial axes). This selection was necessary for simpler statistical analysis but, as indicated by the full topography plots in Figure 5C and supplemental figures S2-S16, our results are robust to different methods of LOT and ROT sensor selection.

### Coding: MI Statistical Significance

We determined the statistical significance of left and right eye coding on LOT and ROT as follows. We compared the maximum MI value within a window of 0 to 400 ms post stimulus, to the 99^th^ percentile over permutations (described above) of the maximum MI over those time points, all electrodes and both features. This resulted in a determination of whether there was any statistically significant coding during that time interval, corrected for multiple comparisons over all electrodes (necessary since we selected LOT and ROT based on MI values), time points and both left and right eye visibility.

### Temporal Precedence: Latency measures

Whereas a human observer can easily determine which of two time varying noisy signals leads the other, it is not straightforward to rigorously define and quantify this. Here, we are interested in the relative timings of the contralateral and ipsi-lateral eye MI on LOT and ROT, rather than their amplitudes. We therefore first normalized the MI curves to their maximum value (in the window 150-250 ms post-stimulus, as plotted in Figure 4A and B). The variability of these curves makes any latency measure that depends on a specific time point problematic due to a lack of robustness. To illustrate, consider the time courses in Figure 4A and B where a simple peak-to-peak measure would result in a value ~100ms for the left eye (blue curves), and – 30ms for the right eye (red curves). These values do not sensibly reflect the actual relationship – for left eye the value seems too high, and for the right eye it is in the wrong direction; the solid contra-lateral curve appears to lead the dashed ipsi-lateral curve in the period 130-180 ms. Similar problems would affect any measure that compares two specific points.

Hence, we considered the latency of the MI curves not over a single value, but over a range of y-axis values (Figure 4C). We restricted our analysis to 100-300 ms post-stimulus, and considered normalized MI values where both curves were significant. We split the range of normalized MI values above the highest significance threshold into 100 values and calculated the mean latency between the two curves over these values (indicated by grey lines in Figure 4C). This is equivalent to integrating the latency of the curves over the y-axis and normalizing by the y-axis range.

### Coding Equivalence: Redundancy

Redundancy (equivalent to negative interaction information (Cover and Thomas, 1991)) quantifies the MI overlap between two variables—that is the amount of MI about eye visibility that is common to both EEG responses. We calculated this as described in the main text and illustrated in Figure 5B. As previously noted, all MI values were calculated with CMI, conditioning out (i.e. eliminating) the contribution of the alternative eye feature:

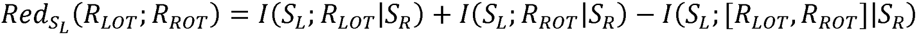

*R*_*LOT*_/*R*_*ROT*_ represent the response across trials at each electrode at the appropriate MI peak (here left eye). Redundancy is bounded above by each of the individual MI values and the MI between the two EEG responses (Cover and Thomas, 1991). We therefore normalized by the minimum of these three quantities:

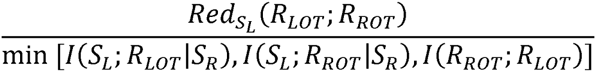

We calculated redundancy between LOT and ROT for each eye feature at the time of the peak of the MI timecourse (within 50-250ms; Table 1).

**Table 1:** Time of MI peak used for redundancy calculation. Median (min max) in ms.

|  | Left eye | Right Eye |
| --- | --- | --- |
| **ROT** | 167 (140 188) | 180 (156 246) |
| **LOT** | 177 (148 216) | 162 (140 180) |

For the redundancy topography plots in Figure 5C and Supplemental Figures S2-S16 we fixed the contra-lateral electrode and MI peak time as above (i.e. ROT for left eye, LOT for right eye) as a seed response. For each eye, we then calculated the redundancy between this seed response and every other sensor and time point where there was significant MI about that eye (p=0.01 with multiple comparison correction as described above).

We also computed MI directly between *R*_*LOT*_ and *R*_*ROT*_ conditioning out any variation due to either stimulus: (*R*_*ROT*_;*R*_*LOT*_|[*S*_*R*_,*S*_*L*_]). Since it is not possible to define a permutation scheme to for multivariate conditional mutual information we performed 1000 bootstrap samples (resampling with replacement), and took the 0.5^th^ percentile as the lower bound of a 99% confidence interval.

## Acknowledgements

We thank Cesare Magri and Daniel Chicharro for useful discussions. KJ is supported by a BBSRC DTP (WestBio) Studentship. JG and PGS are supported by the Wellcome Trust [098433, 107802]. GAR, PGS and NJvR are supported by the BBSRC [BB/J018929/1]. SP is supported by the VISUALISE project of the Future and Emerging Technologies (FET) Programme within the Seventh Framework Programme for Research of the European Commission (FP7-ICT-2011.9.11) under grant agreement number FP7-600954.

## Supplemental Figures

**Figure S1.**
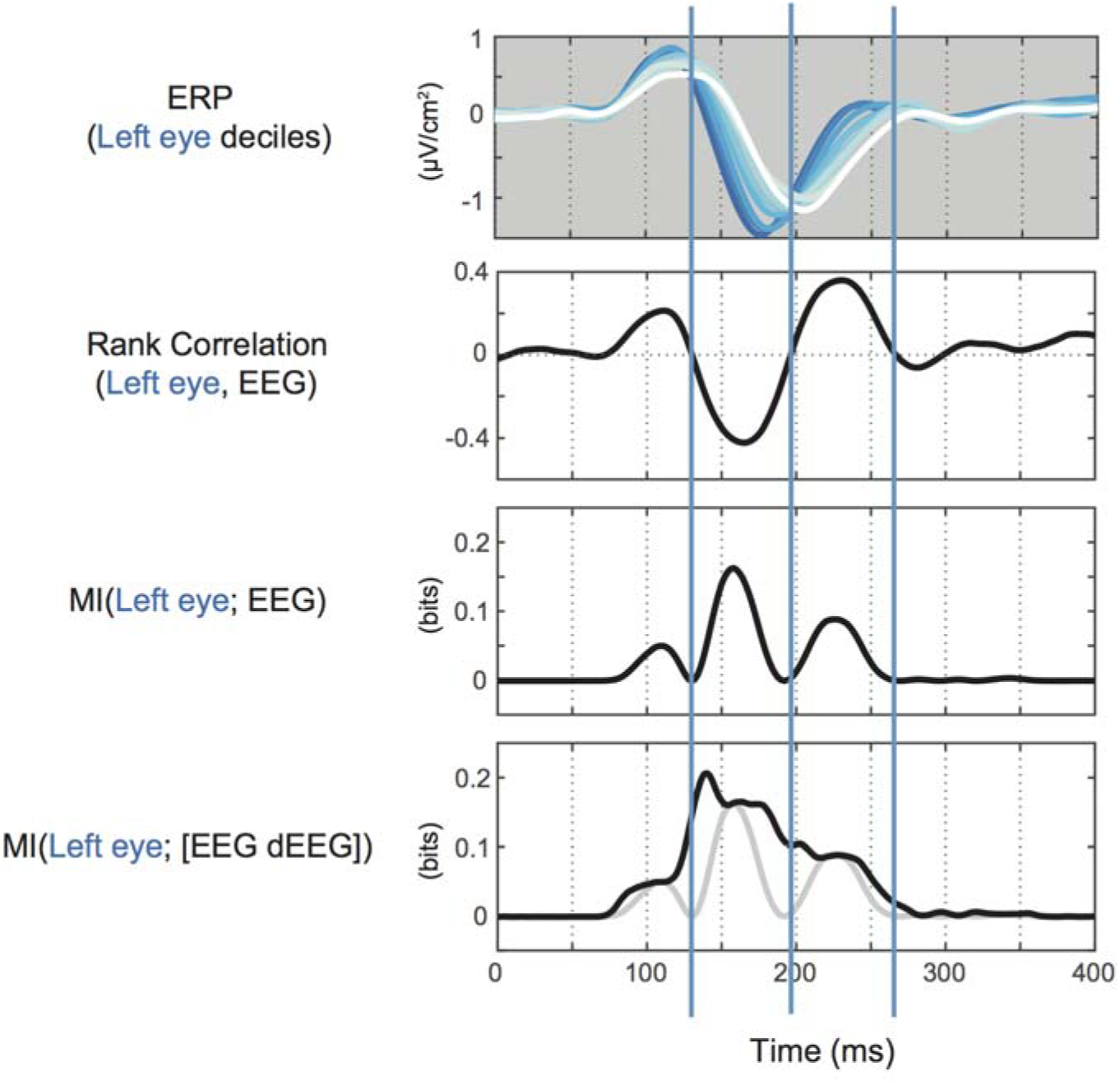
*Mutual information calculated on bivariate EEG response including temporal gradient.* From top to bottom, plots show (i) the ERP calculated from trials in each decile of left eye visibility (c.f. Figure 3) on ROT of observer 1, (ii) Spearman’s rank correlation between the left eye visibility and EEG voltage, calculated separately for each time point, (iii) MI between the left eye visibility and EEG voltage calculated separately for each time point and (iv) MI between left eye visibility and the 2D EEG response consisting of voltage and temporal gradient, calculated separately for each time point. Zero crossings of rank correlation, where the directionality of the voltage modulation changes, are indicated with vertical bars.

**Figure S2.**
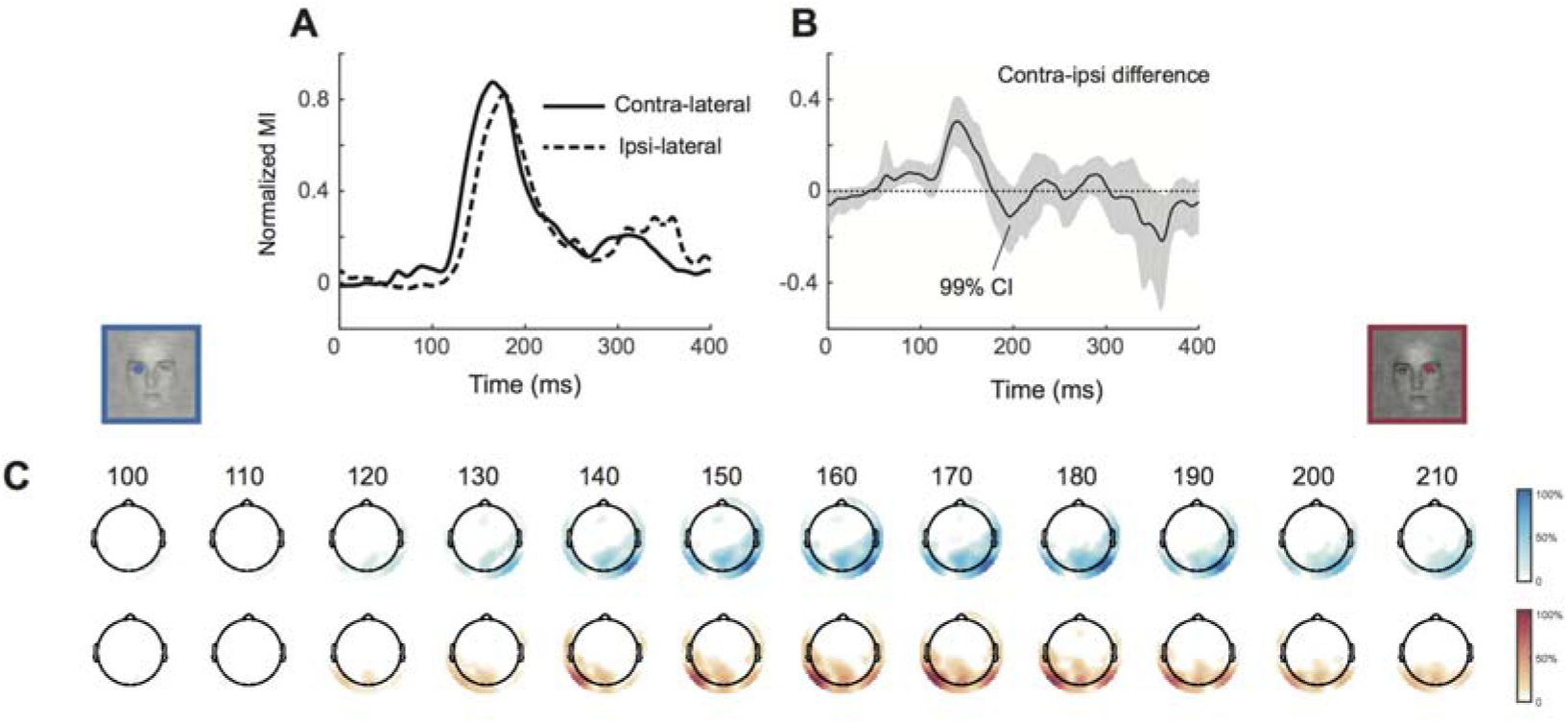
*Group average results for temporal precedence and coding equivalence.* **A.** Mean normalized MI time course over all instances with bilateral coding significance (observer and left or right eye, N=26 / 30) for contra‐ and ipsi-lateral sensors (LOT/ROT). **B.** Mean contra-ipsi difference over instances, with 99% bootstrap confidence interval. **C.** Mean normalized redundancy topography sequences calculated as described in the text.

**Figures S3-S16.**
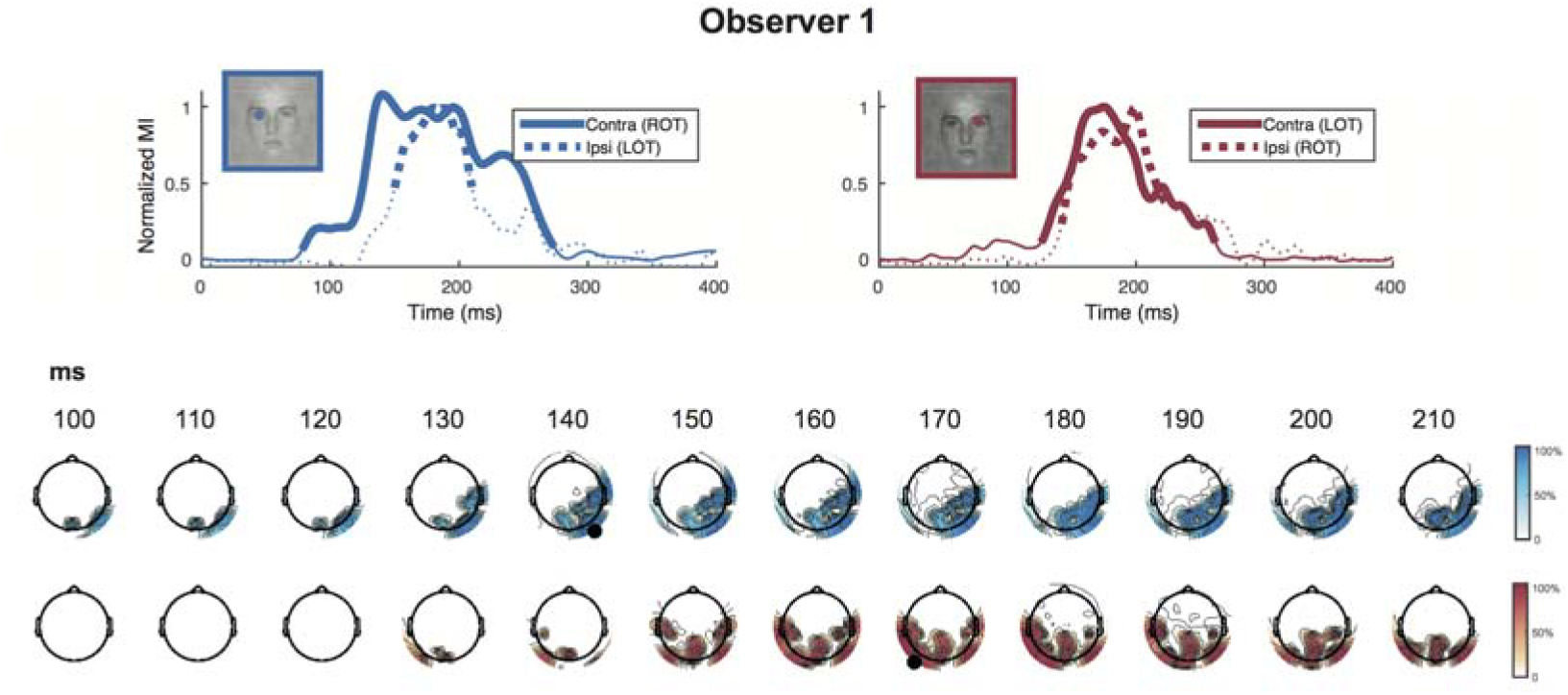
*Individual observer results for temporal precedence and coding equivalence.* Time courses show the normalized MI time courses for the left eye visibility (blue, left plot) and right eye visibility (red, right plot) on the contra-lateral (solid line) and ipsi-lateral sensors (dashed line). Topography sequences show normalized redundancy (seed sensor and time indicated with black circle).

**Figures S4.**
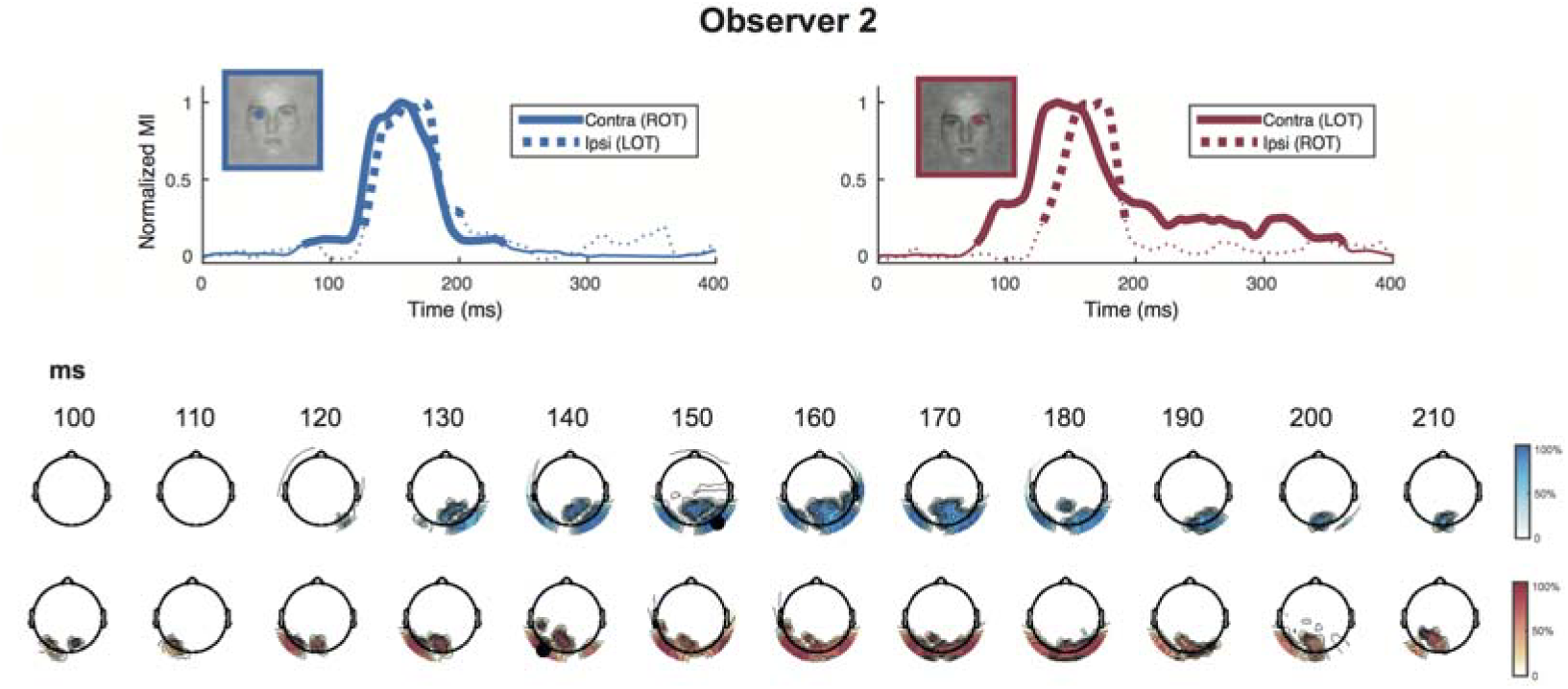

**Figures S5.**
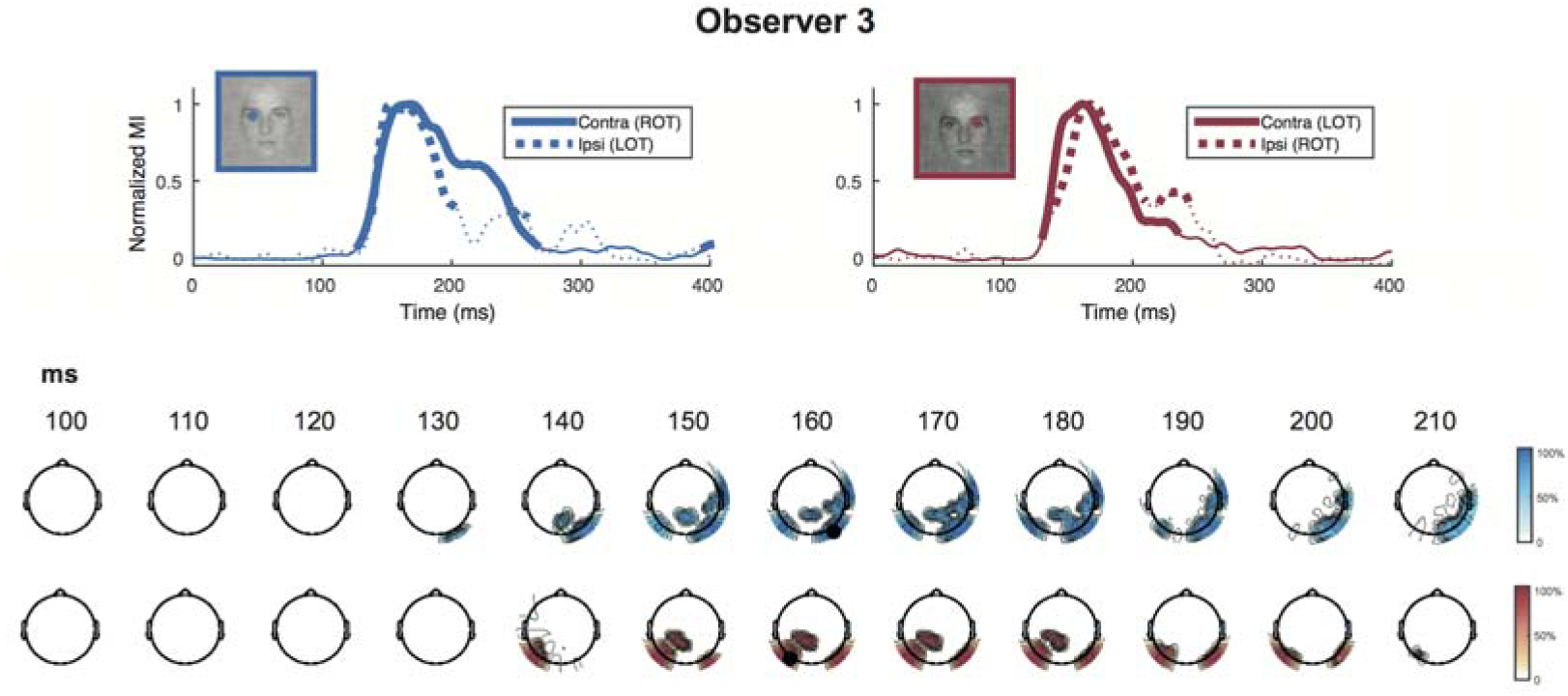

**Figures S6.**
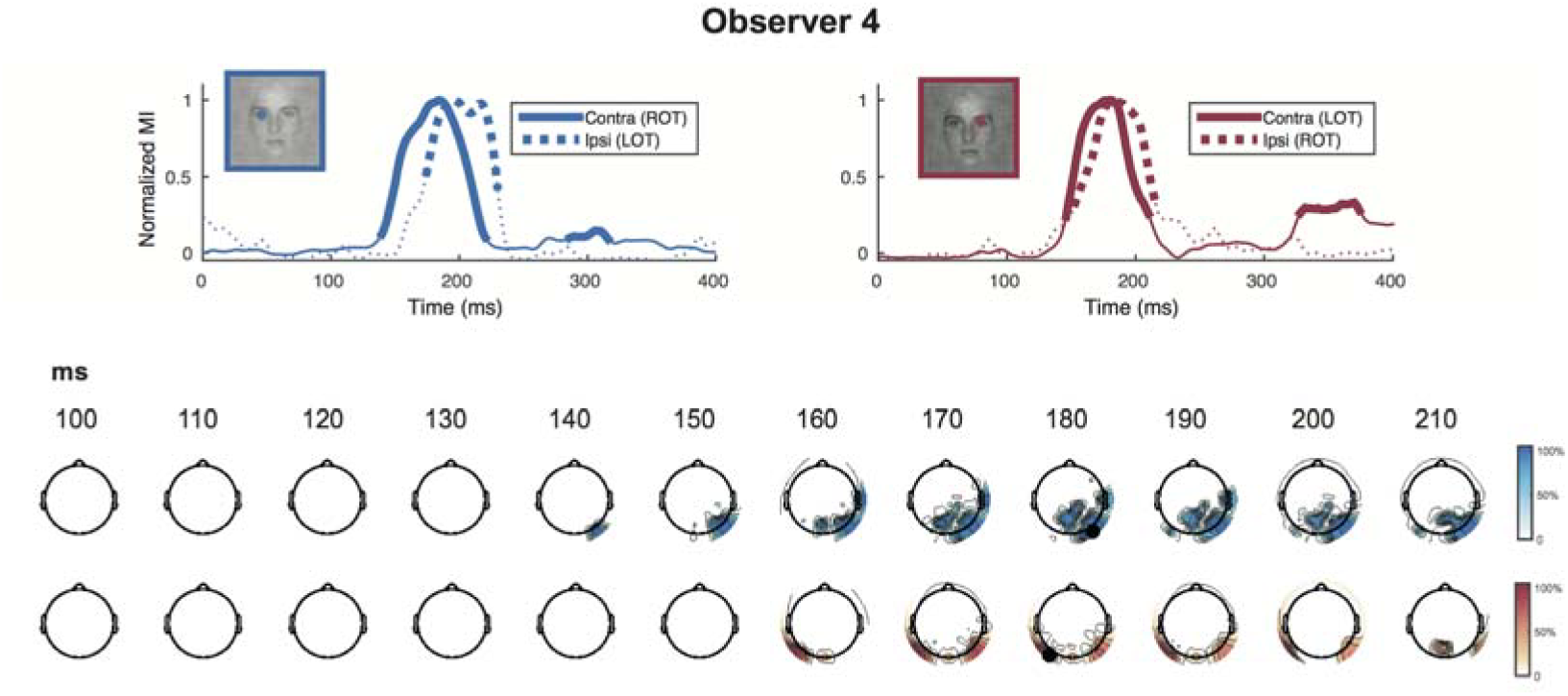

**Figures S7.**
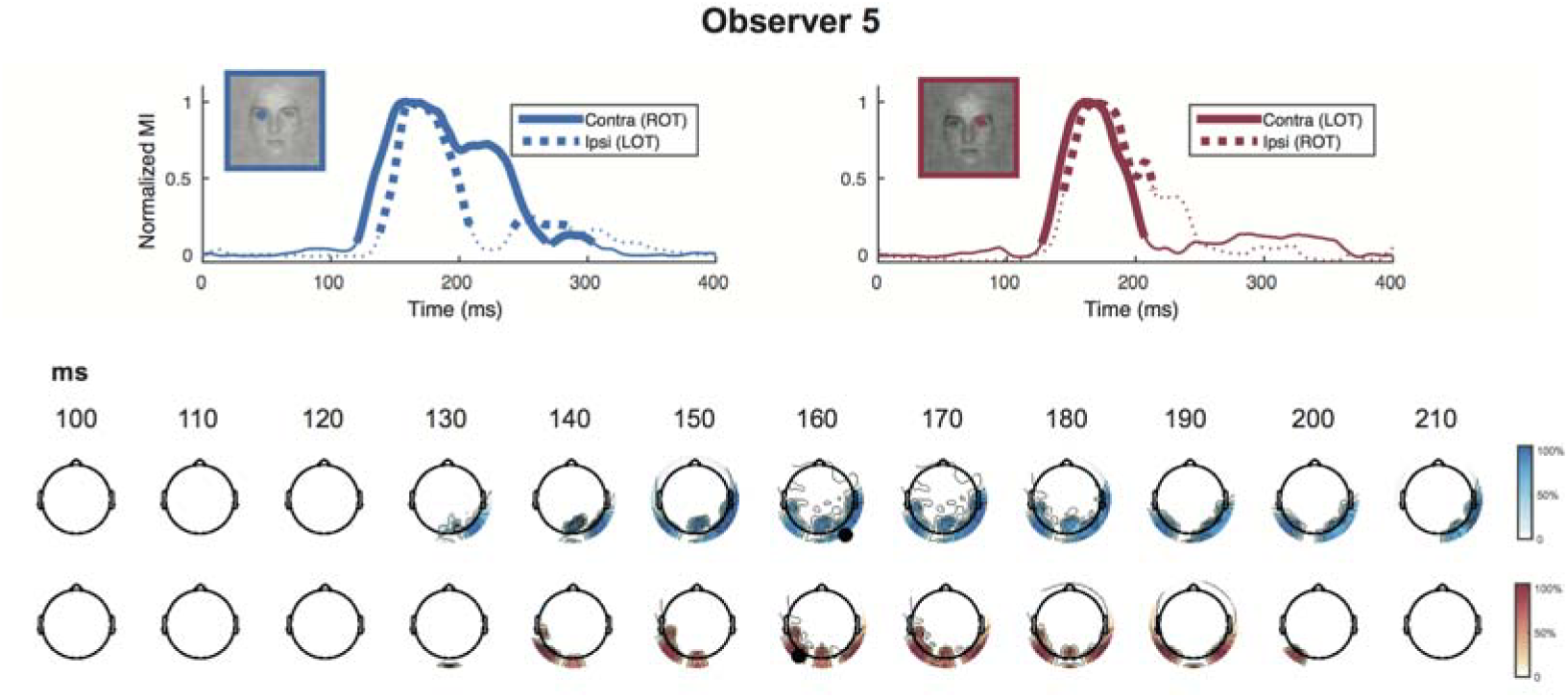

**Figures S8.**
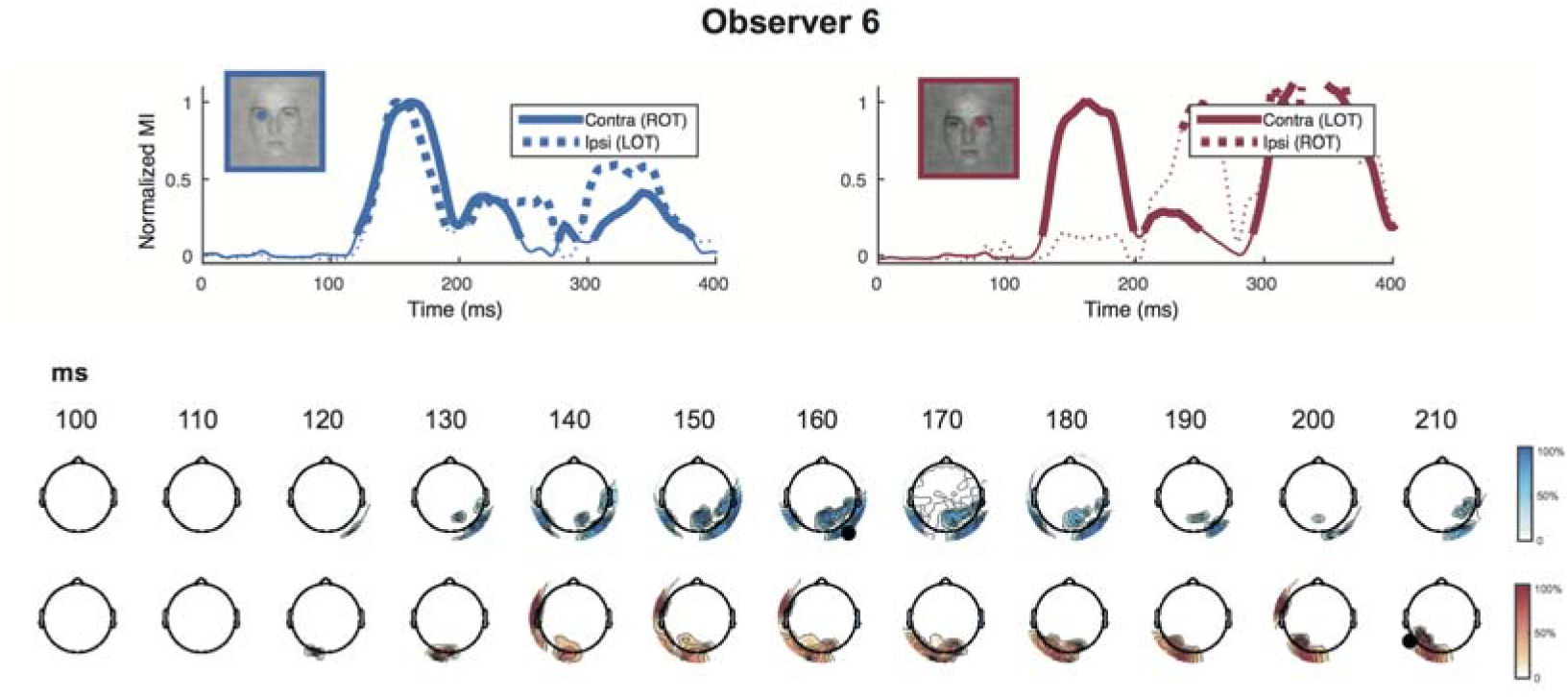

**Figures S9.**
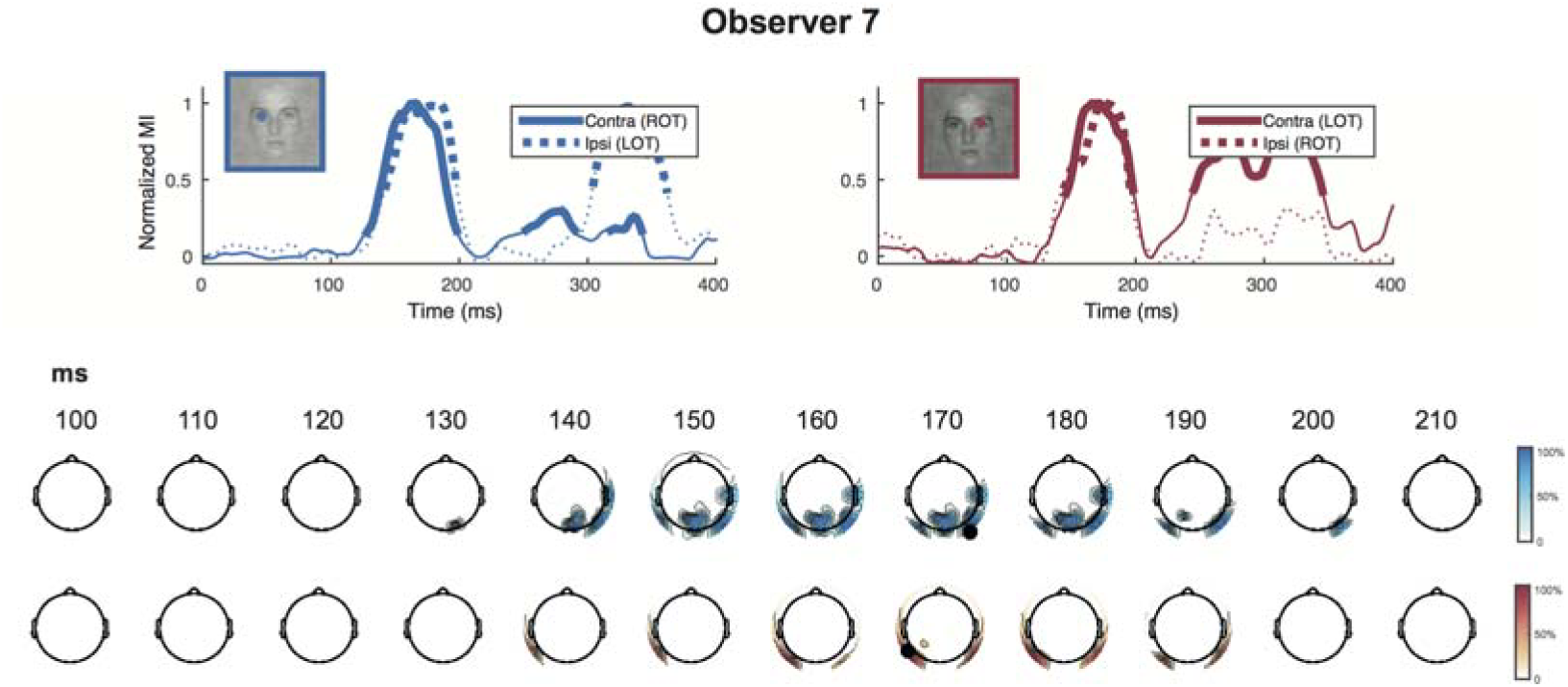

**Figures S10.**
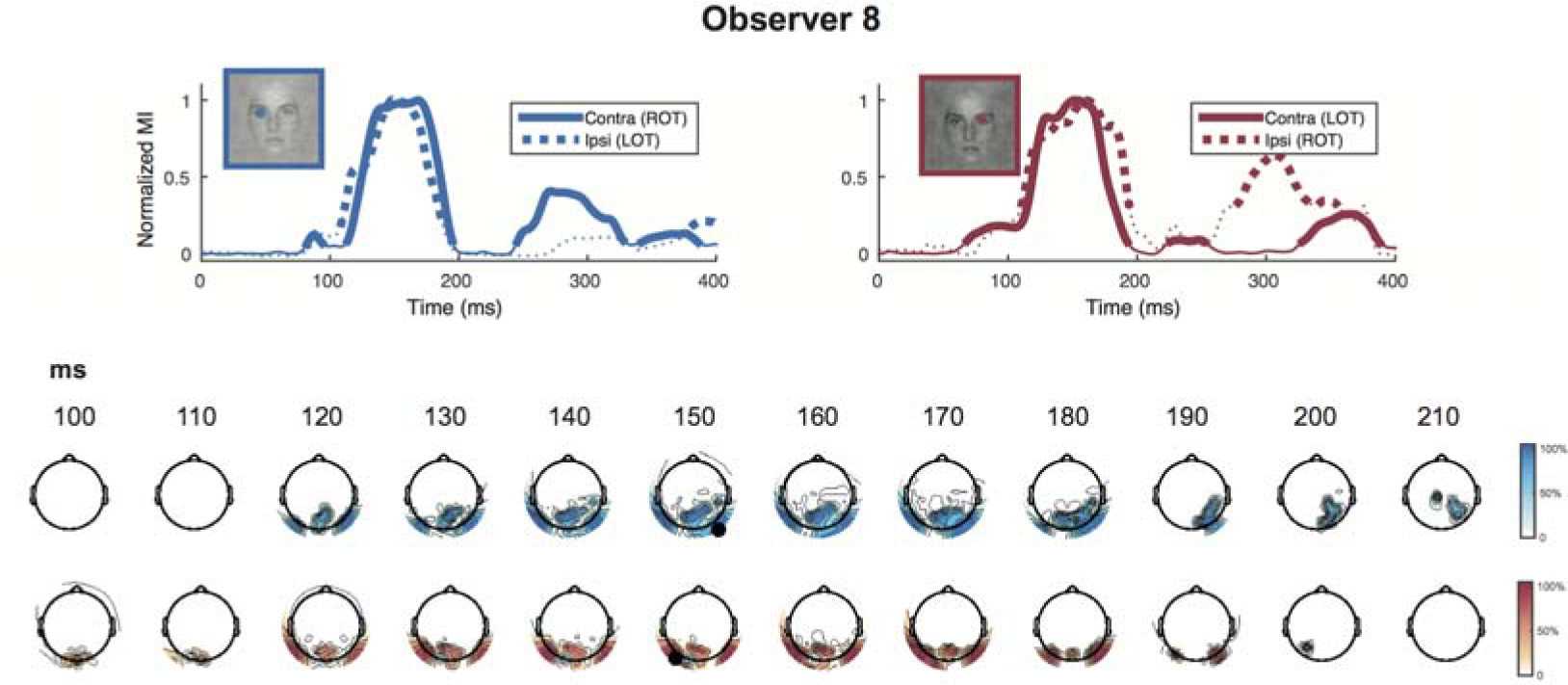

**Figures S11.**
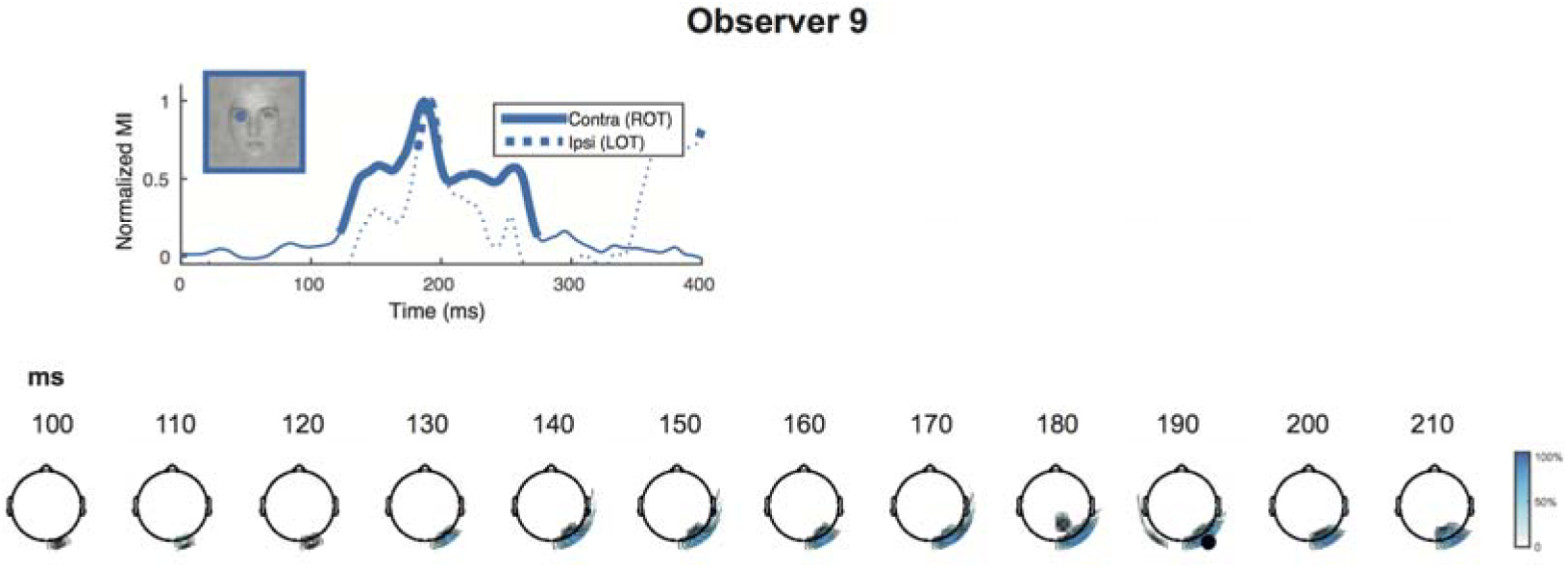

**Figures S12.**
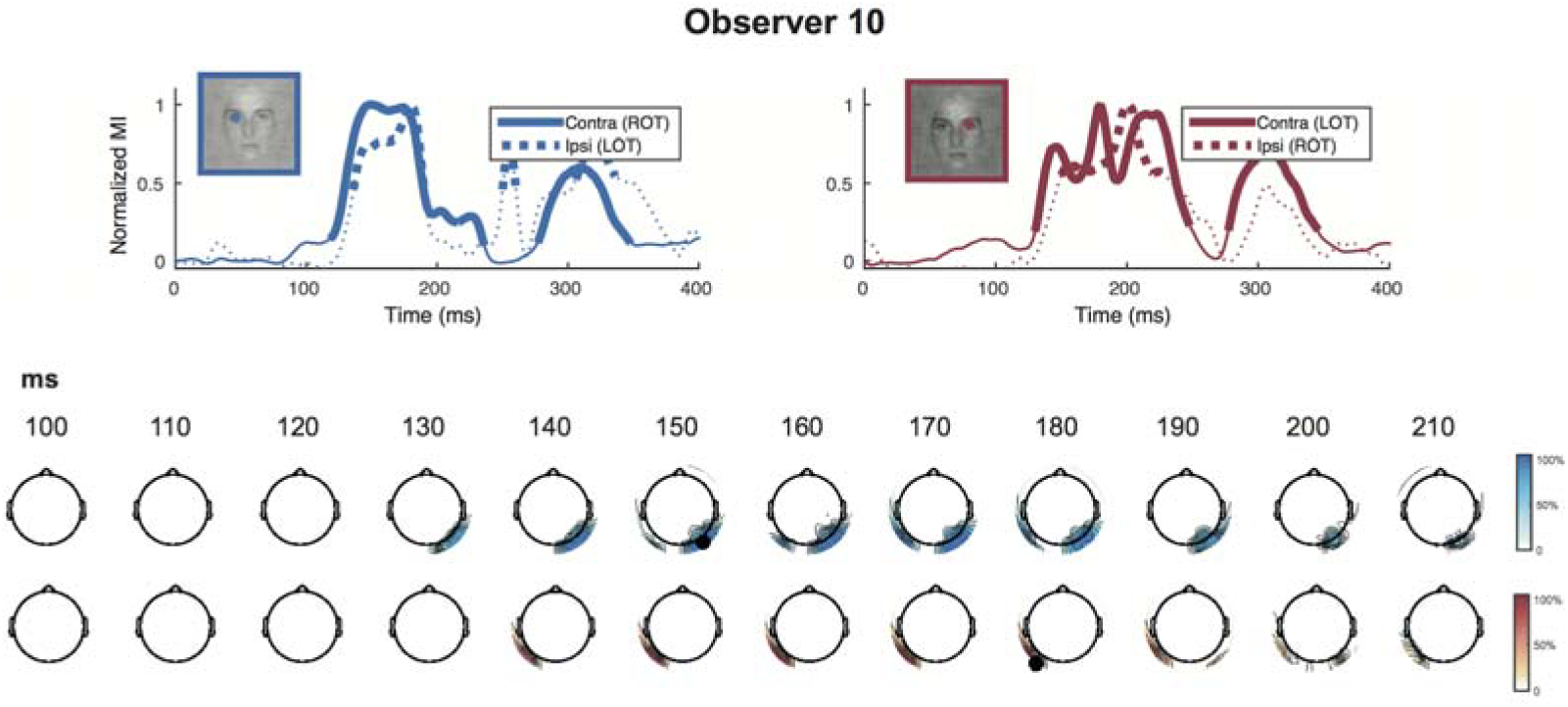

**Figures S13.**
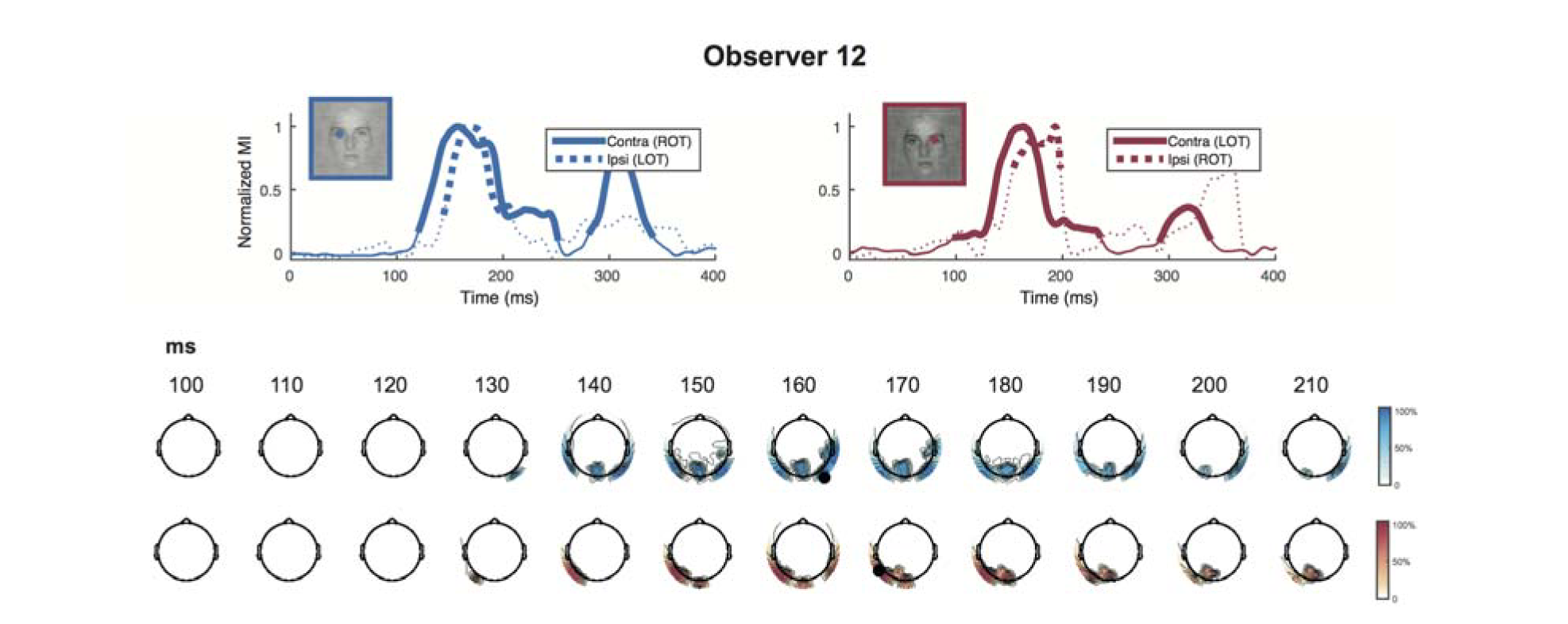

**Figures S14.**
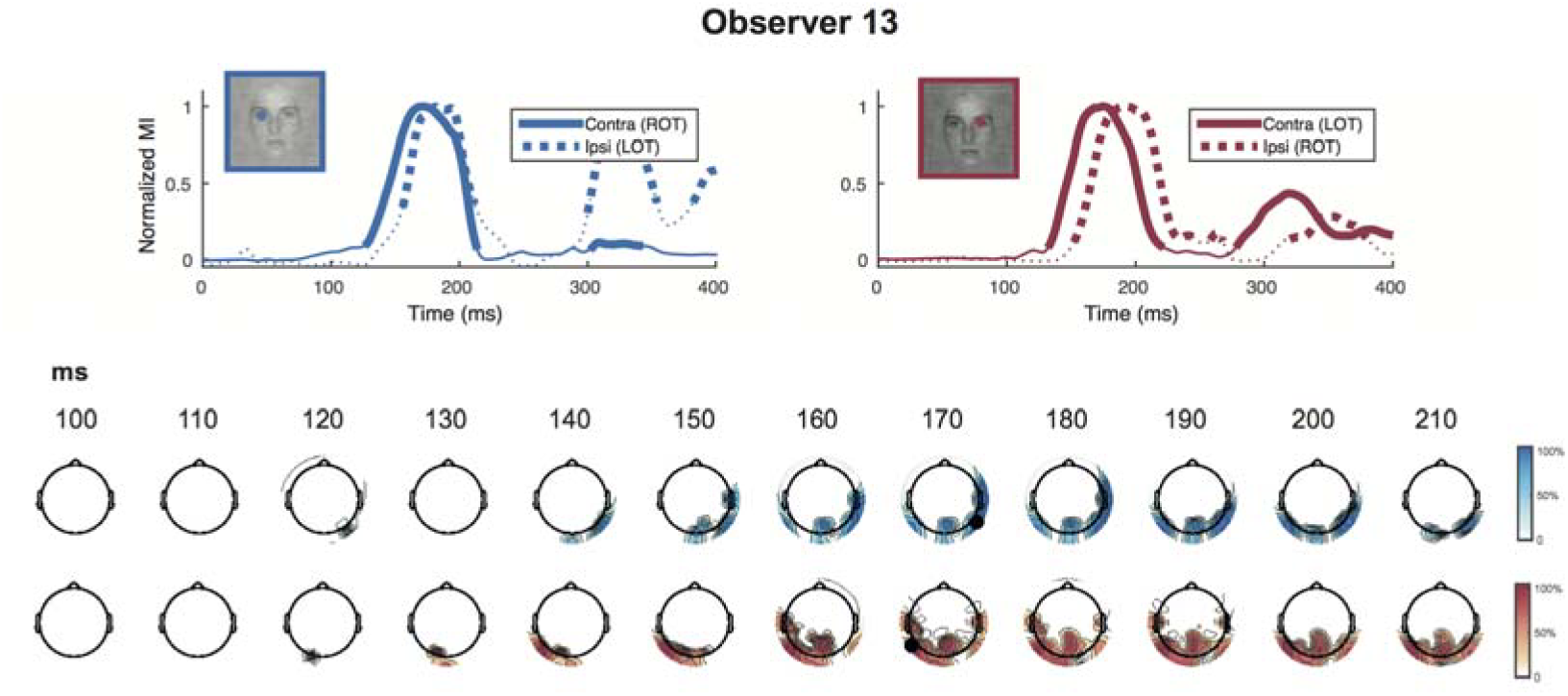

**Figures S15.**
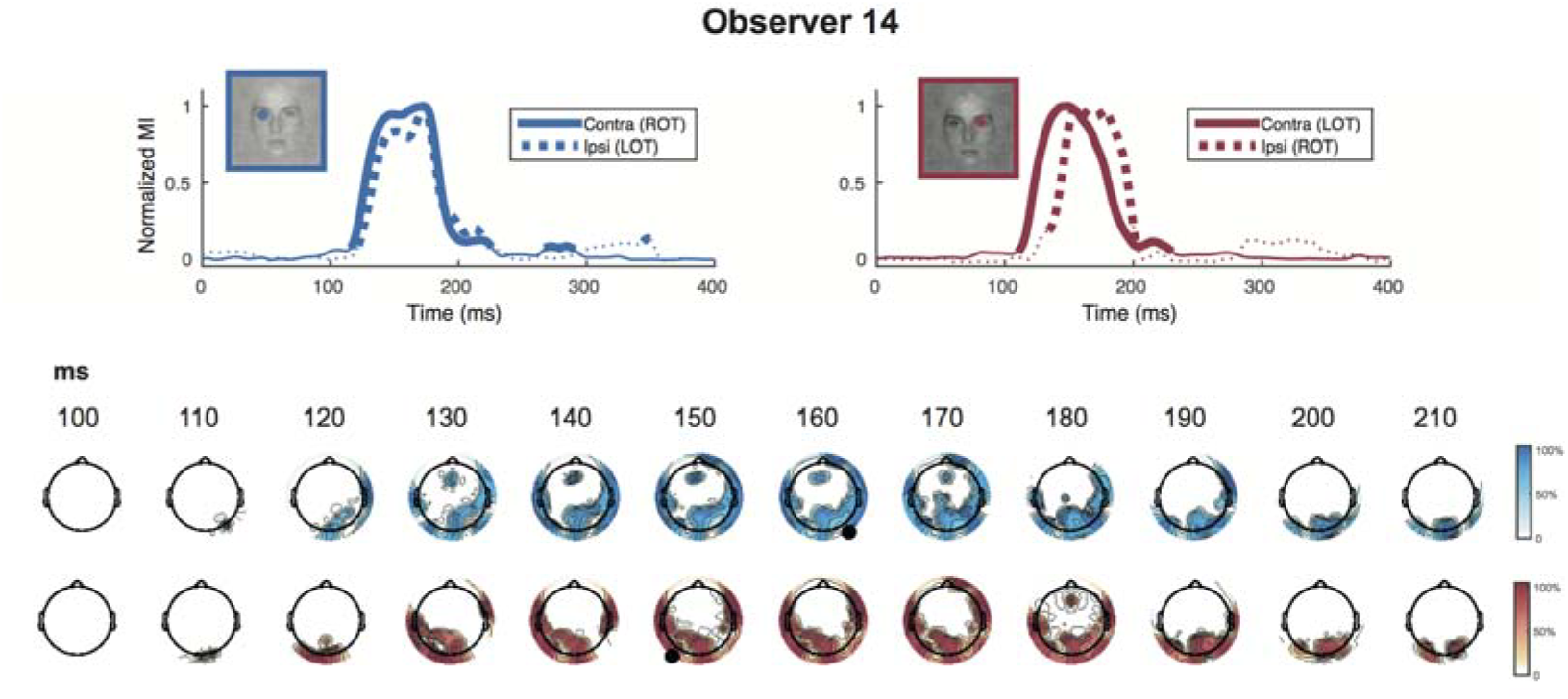

**Figures S16.**
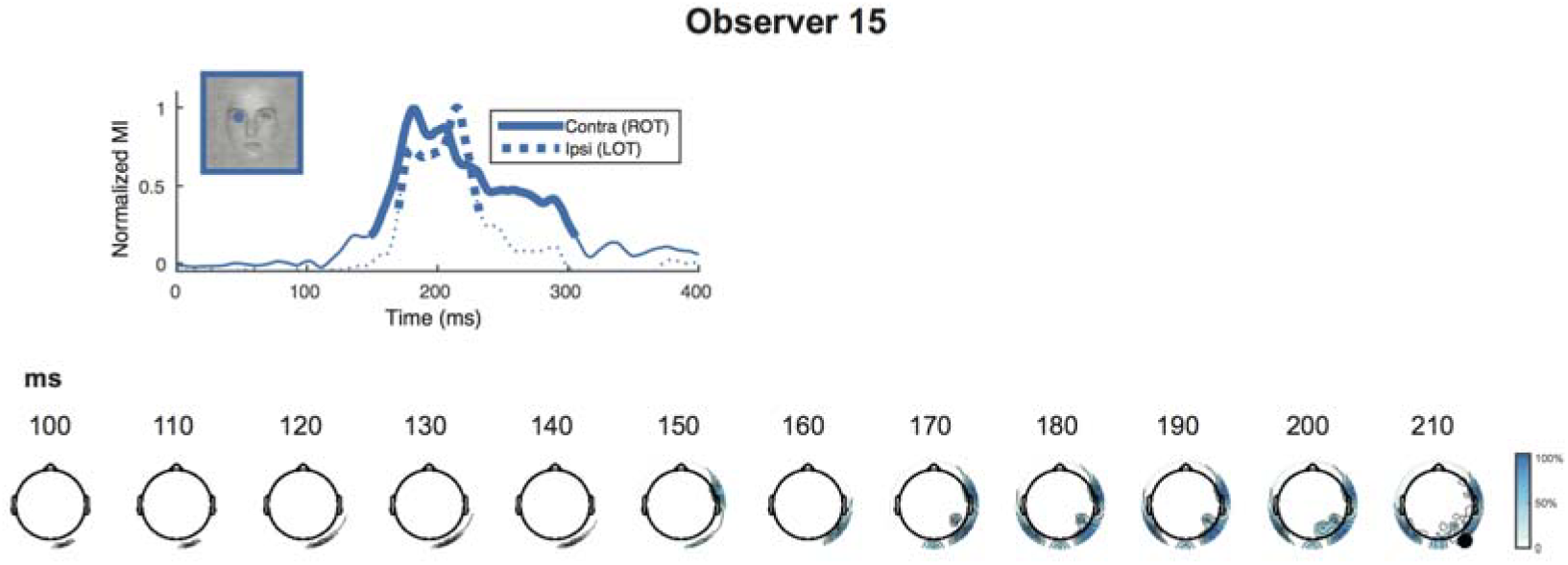

